# Structural basis for tetraspanin-dependent surface export and adhesive function of integrin α3β1

**DOI:** 10.64898/2026.09.01.748654

**Authors:** Lauren Litzau, Elizabeth K. Wren, Andrew C. Kruse, Stephen C. Blacklow

## Abstract

Integrin α3β1 (ITGα3β1) is a member of an integrin subfamily that binds to laminin proteins and promotes attachment of epithelial cells to the basement membrane. ITGα3β1 forms a complex with the tetraspanin CD151, and loss-of-function mutations in both ITGα3 and CD151 cause epidermolysis bullosa, a severe skin blistering disease resulting from a defect in basement membrane attachment. Here, we report the cryoEM structure of an ITGα3β1 complex with CD151 and show that mutation of CD151 at the binding interface disrupts complex formation in cells. Strikingly, CRISPR-mediated knockout of CD151 leads to a variably penetrant ITGα3β1 surface export defect that is restored by re-expression of wild-type but not interface-mutated CD151. Together, these studies define the molecular basis for binding of CD151 to ITGα3β1, and show that CD151 promotes ITGα3β1 surface export, providing a biochemical explanation for the CD151 loss-of-function phenotype in epidermolysis bullosa.

## Introduction

Integrins are heterodimeric receptors of noncovalently associated α and β subunits that link cells to components of the extracellular matrix and to other cells. They respond to changes in the extracellular environment and organize the intracellular cytoskeleton, playing critical roles in cell adhesion, migration, cellular organization and development.^1^ Both integrin subunits are type I transmembrane glycoproteins with a large extracellular domain and a small intracellular tail. In humans, there are 24 known heterodimeric α/β pairings with distinct ligand specificities for fibronectin/RGD ligands, collagen, laminin, or leukocyte antigens.

Tetraspanins are four-pass transmembrane domain proteins with diverse biological roles influencing trafficking, membrane localization, and signaling of their partner proteins.^2^ In humans, 10 of the 33 tetraspanin family members can interact with integrins, but the molecular details of how tetraspanins influence integrin function are not well understood.^3^

Among tetraspanin proteins, CD151 stands out for its ability to associate with all laminin binding integrins, binding most stably to the laminin-binding integrin α3β1 (ITGα3β1).^4^ Silencing mutations in either CD151 or ITGα3 result in epidermolysis bullosa, a skin blistering condition consistent with impaired laminin binding and basement membrane attachment. Kidney defects are observed in both CD151 and ITGα3 knockout (KO) mice (*11*), and in cancer, CD151 and ITGα3β1 are commonly upregulated together in various epithelial tumors, with high CD151 expression correlating with poor prognosis and metastasis (*12*). These phenotypic similarities strongly suggest that CD151 loss-of-function is associated with diminished integrin activity, yet the molecular basis for this effect remains unclear.

Here, we used a combination of structural studies of the ITGα3β1-CD151 complex and cell-based assays to determine the molecular basis for functional dependence of ITGα3β1 on CD151. Using single-particle cryoEM, we found that CD151 binds to a conserved surface on integrin α3 opposite the β1 interface at a site predicted to be accessible independent of integrin conformation. In CD151 knockout cells, there is an ITGα3β1 export defect of variable penetrance that is suppressed by expression of wild-type CD151 but not by ITGα3β1 binding-defective CD151 mutants. Together, these findings reveal the molecular basis for integrin binding by a tetraspanin protein and identify a biochemical rationale for the overlapping phenotypes of patients with loss-of-function mutations in CD151 and integrin α3 in epidermolysis bullosa.

## Results

### Cryo-EM structure of ITGα3β1-CD151

To produce samples of a ITGα3β1-CD151 complex suitable for data collection by cryoEM, we used a protein fusion approach^5^, in which we connected the C-terminus of full-length ITGα3 to the N-terminus of full-length CD151 using a flexible GGSx4 linker (Figure S1, related to Figure 1). This strategy maintained topology, increased expression, and ensured a 1:1 stoichiometry of ITGα3 to CD151. We purified the complex using size-exclusion chromatography (Figure S1C, related to Figure 1), collected cryoEM images for single particle analysis, and reconstructed a map at a nominal resolution of 3.8 Å after data processing (Figure 1A and Figure S2, related to Figure 1). The ITGα3β1 headpiece region was well defined in this reconstruction, but the resolution of the membrane proximal and transmembrane regions was poorer and only visible at a lower contour threshold.

**Figure 1.**
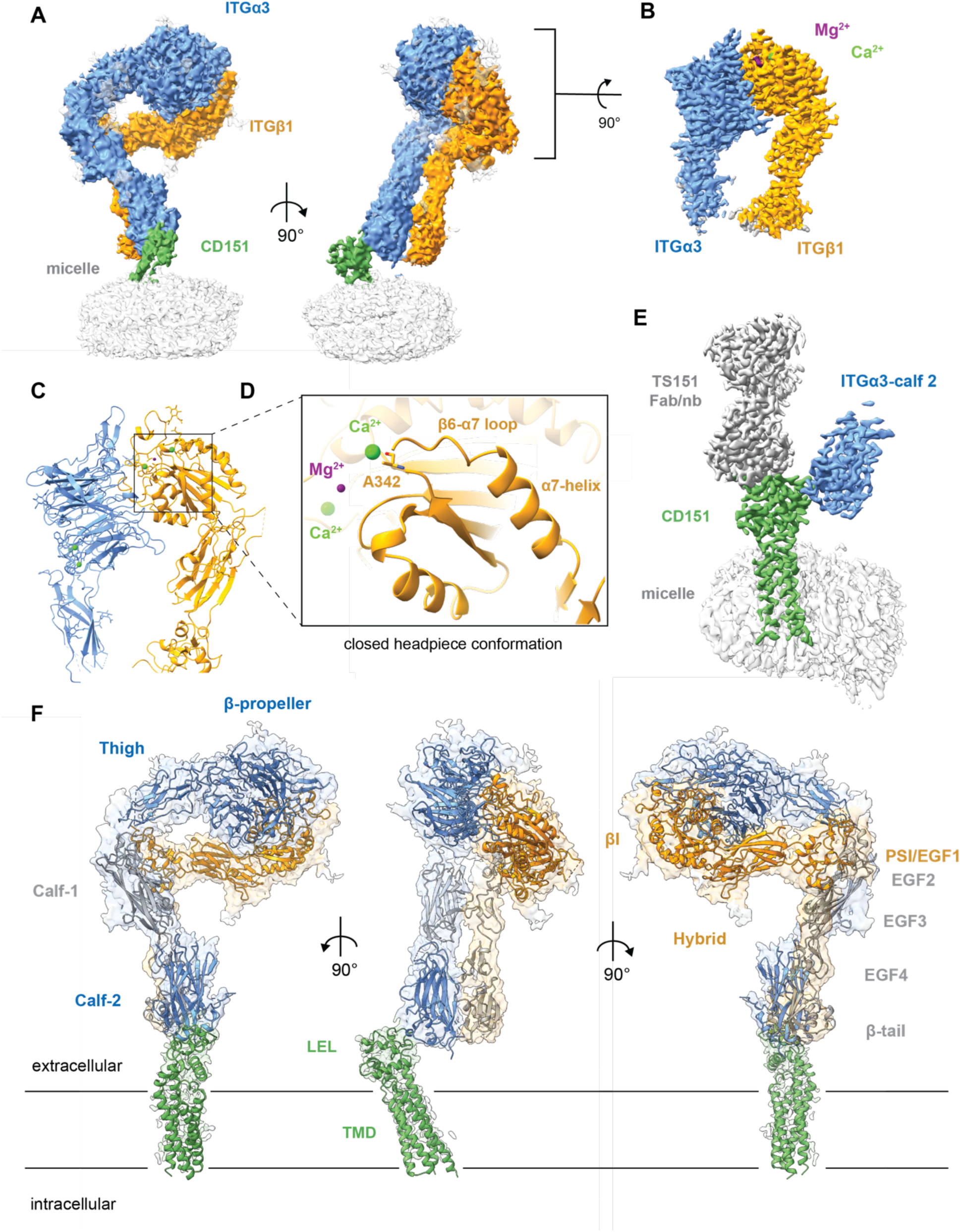
Cryo-EM structure of the ITGα3β1-CD151 complex. (A) Density map of the full length ITGα3β1-CD151 complex. (B) Density map of the ITGα3β1 headpiece after local refinement. (C) Cartoon rendering of the ITGα3β1 headpiece. (D) Zoomed in view of the ITGα3β1 headpiece ADMIDAS site, showing A342 of the ITGβ1 subunit rendered as sticks. (E) Density map of a truncated complex containing the ITGα3 Calf-2 domain fragment (blue), CD151 (green), and the TS151 anti-CD151 Fab bound to an anti-Fab nanobody (gray). (F) Cartoon representation of a composite model of the full-length complex, derived from overlaying the two maps. The coordinates for the headpiece were from the full-length complex after local refinement, the coordinates for the ITGα3 Calf-1 domain, ITGβ1 EGF repeats 2-4 and the ITGβ1 β-tail domain (shown in gray) were from an AlphaFold2 model docked into the density map of the full length complex, and the coordinates for CD151 and the calf-2 domain were from the atomic model built into the density map of the truncated complex shown in (E).

Local refinement of the ITGα3β1 headpiece yielded a 3.1 Å map suitable to build a model of the β-propeller and thigh domains of ITGα3 (residues 33-610), and the βI, hybrid, thigh, and EGF1 domains of ITGβ1 (residues N25-D505) (Figure 1B). This headpiece region is in a bent, closed conformation, consistent with prior work showing that the closed state is typically the predominant species of unliganded integrins in cells.^6^ The ITGβ1 α7 helix is in its closed, low-affinity configuration where the carbonyl of A362 in the β6-α7 loop of ITGβ1 coordinates the Ca^+2^ of the adjacent to the metal ion dependent site, or ADMIDAS (Figure 1C,D).^7^

To acquire a higher resolution view of the CD151-ITGα3 interface, we designed a fusion protein linking a truncated ITGα3 calf-2 domain to CD151 (Figure S1B, related to Figure 1), and complexed it with an anti-CD151 Fab called TS151^8^ and an α-Fab hinge-stabilizing nanobody^9^ to increase particle mass (Figure S1D, related to Figure 1). We determined the structure of this assembly at an overall resolution of 2.9 Å (Figure 1E). By superposing the map of the calf-2 domain of this truncated complex onto the map of the calf-2 domain in the full-length complex, it was then possible to construct a composite model of the complex (Figure 1F), using an Alphafold3 model for the calf-1 domain of the α3 subunit, and EGF-like domains 2-4 and the β-tail domain of the β1 subunit. It was not possible, however, to identify density corresponding to the transmembrane or cytoplasmic regions of either integrin subunit in the maps.

### CD151 binds ITGα3 through an extracellular site

Contacts between CD151 and the integrin are exclusively between a basic surface on the large extracellular loop of CD151 and the calf-2 domain of ITGα3, independent of the ITGβ1 subunit (Figure 2A). The interface has a total buried surface area of only 955 Å^2^ and is dominated by salt bridge and hydrogen bond interactions between a basic surface on CD151 and an acidic patch on the calf-2 domain of ITGα3 (Figures 2B-D). The contact surface is centered on H205 of CD151, which forms electrostatic contacts with both E782 and E823 of ITGα3. Flanking interactions between K194 of CD151 with E940 of ITGα3, and between R203 of CD151 and E782 of ITGα3, also contribute to this electrostatic complementarity. The direct contact between R203 and E782 is consistent with prior biochemical studies, which reported that mutation of the three-residue QRD sequence that includes R203 to INF disrupts co-immunoprecipitation of ITGα3 with CD151^10^, although our structure shows that only R203 interacts directly with ITGα3 (Figure 2C and Figure S4, related to Figure 2). Additional contacts between CD151 and ITGα3 include contacts from the side chains of T195 and S207 to the ITGα3 backbone, and a side chain interaction between N208 of CD151 and S783 of ITGα3 (Figure S4, related to Figure 2). Although AlphaFold multimer predicted an additional salt bridge interaction between E213 of CD151 and K828 of ITGα3, these residues are too distant to make direct contact in our experimentally determined structure (Figure 2E).

**Figure 2.**
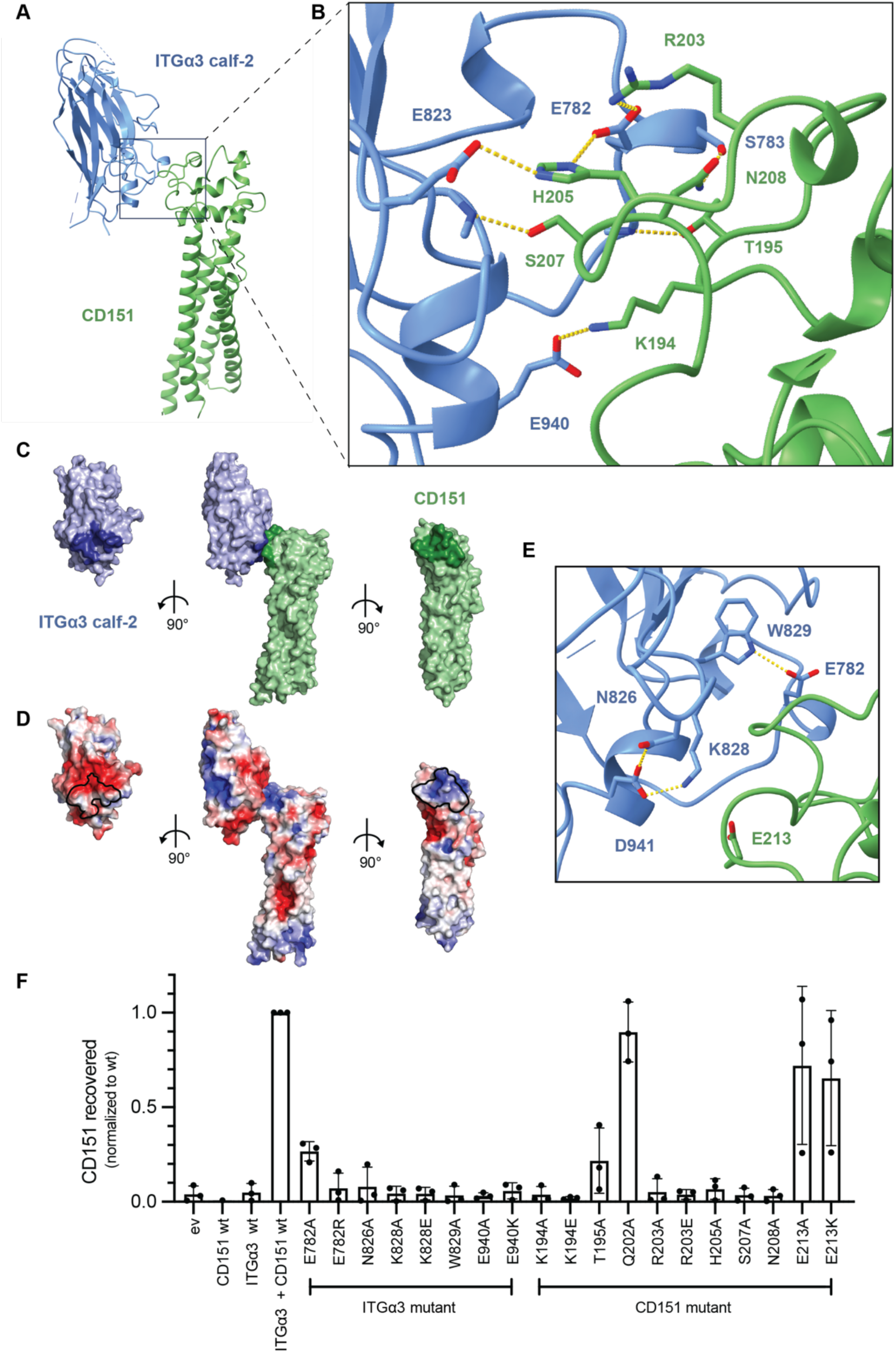
ITGα3 and CD151 interact through an extracellular site. (A) Cartoon representation of the complex between CD151 and the ITGα3 calf-2 domain. ITGα3 is blue, and CD151 is green. (B) Zoomed in view of the contact interface between ITGα3 calf-2 and CD151. Key interacting side chains are shown in stick format. Hydrogen bonds are indicated with dashed yellow lines. (C) Contact interface analysis. Surface representation of the complex (middle) alongside open book views of the ITGα3 calf-2 domain (left) and CD151 (right), colored as in panel (A) with residues at the contact interface colored a darker shade. (D) Electrostatic surface potential at the binding interface. Surface representation of the complex (middle) alongside open book views of the ITGα3 calf-2 domain (left) and CD151 (right), colored by electrostatic potential on a sliding scale from red (acidic) to blue (basic). (E) Cartoon view highlighting ITGα3 residues proximal to the binding interface that stabilize the conformation of the CD151-interacting loop. (F) Effect of mutations in ITGα3 or CD151 on co-immunoprecipitation of these two subunits of the complex. SVG-A TKO cells were transfected with CD151 and with ITGα3 containing a protein C epitope tag. Complexes were immunoprecipitated with an anti-protein C antibody, and the amount of CD1551 recovered was quantified by Western blot by normalizing to the amount of CD151 protein in the input sample before immunoprecipitation (see Figure S6, related to Figure 2). Data are plotted as the mean ± SEM from three biological replicates (n=3).

To evaluate the importance of these contacts and of interface-adjacent residues in complex stabilization and in functional assays, we used CRISPR/Cas9 for genome editing of SVG-A fetal astrocyte cells, which express substantial amounts of CD151, ITGα3, and ITGα6. We engineered SVG-A cells lacking CD151 (see below), ITGα3, or ITGα6, then constructed an ITGα3/ITGα6 double knockout, and finally a triple knockout (TKO) lacking all three proteins (Figure S5, related to Figures 2-4). We transfected the TKO cells with wild-type or mutated forms of CD151 and ITGα3 to test for complex formation in an immunoprecipitation assay, using a protein C affinity tag appended to the C-terminus of ITGα3 (Figure 2F). The immunoprecipitation assay showed that mutations of key ITGα3-contacting residues on CD151, including K194E, T195A, R203A, R203E, H205A, S207A, and N208A, disrupt immunoprecipitation by ITGα3, whereas the Q202A, E213A, and E213K mutations, which are CD151 residues outside of the contact interface, do not interfere with immunoprecipitation by ITGα3 (Figure S6, related to Figure 2). On the other side of the interface, introducing mutations into ITGα3 on the acidic patch at E940 and E782 either partially or complete interfere with immunoprecipitation of CD151, as do mutations of W829, N826, and K828, which form hydrogen bond or salt bridge interactions between ITGα3 loops and are proximal to the CD151-ITGα3 interface, but do not directly interact with CD151 (Figure S6, related to Figure 2). The ITGα3 residues required for stable complex formation with CD151 are conserved among the three laminin-binding integrin alpha subunits. The only divergent position is serine 783, which is on the rim of the interface, and replaced by glutamine in ITGα6 and arginine in ITGα7. These substitutions can be readily accommodated without disrupting the interface, consistent with published reports that CD151 co-immunoprecipitated with all laminin binding integrins.^11^

### CD151 KO diminishes ITGα3β1 surface export

To assess the effect of CD151 on cell surface delivery of ITGα3β1, we compared the surface staining of wild-type and CD151 SVG-A knockout cells. After cells were transfected with a gRNA/CAS9 targeting plasmid also expressing GFP, single cells were sorted for GFP positivity using flow cytometry. Knockout clones were identified by the loss of reactivity with the anti-CD151 antibody TS151 on Western blot and confirmed by sequencing genomic DNA from the targeted locus after PCR amplification.

We compared the amount of ITGα3 surface staining in individual CD151 knockout clones with that of individual parental cell clones that remained CD151 wild-type after receiving the targeting vector. Although there was heterogeneity in the magnitude of the effect, there was a substantial reduction in ITGα3 surface staining in the CD151 knockout clones that was statistically significant when compared to that in parental, CRISPR-treated wild-type clones (Figure 3A, p<0.0001, unpaired t-test). After transfection of wild-type CD151 into the knockout cells, the cell-surface staining of ITGα3 was restored to parental cell levels within the CD151 expressing population (Figure 3B), regardless of whether the CD151 knockout clone was highly, moderately, or weakly defective in ITGα3 surface staining (Figure 3C). Experiments using CD151 mutant proteins showed that K194E and H205E were most ineffective at rescuing ITGα3 surface staining, that alanine substitutions K194A, R203A, R203E, H205A, S207A, and N208A, along with the QRD to AAA mutation, had an intermediate rescue phenotype, and that Q202A, D204A, E213A, and E213K were as effective as wild-type CD151 in rescuing ITGα3 surface staining (Figure 3D). When two charge reversal mutations of CD151 were combined in the K194E/R203E, K194E/H205E, and R203E/H205E proteins, or when all of these sites were mutated to glutamate in the K194E/R203E/H205E triple mutant, the protein failed to increase ITGα3 surface staining above the basal amount seen in the knockout clone. These findings are highly consistent with results of the immunoprecipitation studies and with the features of the CD151-ITGα3 interface seen in the structure of the complex, in which the critical contacts are electrostatic interactions of the acidic surface of ITGα3 with the basic patch on CD151 centered around H205, supplemented by additional contacts to K194 and R203.

**Figure 3.**
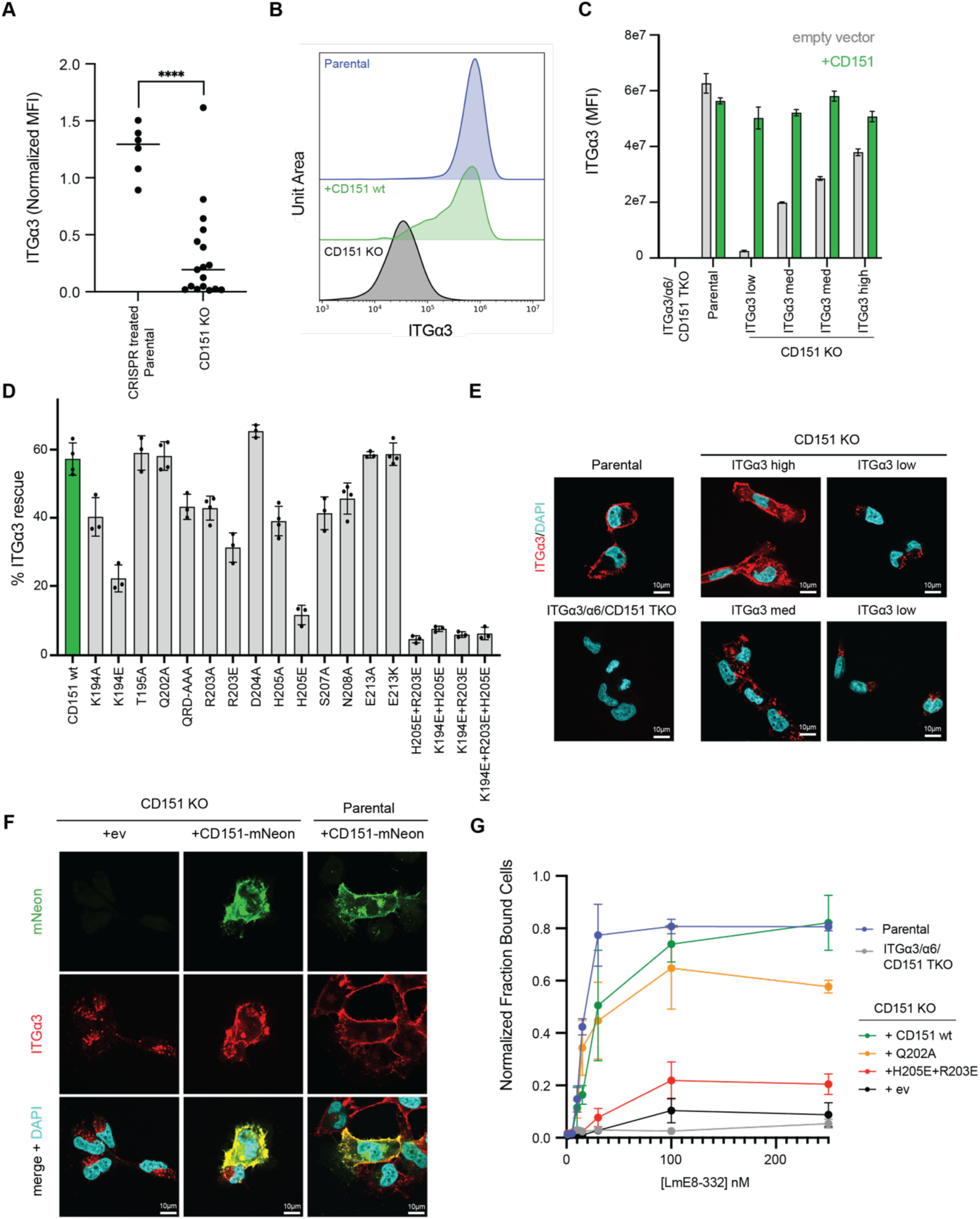
CD151 KO diminishes surface export and function of ITGα3β1. (A) Plot comparing ITGα3 surface staining (Mean fluorescent intensity; MFI) of CRISPR-treated parental cell clones with CD151 KO clones, normalized to ITGα3 surface staining in untreated SVG-A cells. Each dot represents a single cell CD151 KO or CRISPR treated parental clone. Statistical significance was determined by an unpaired t-test, ****p<0.0001 (n ≥ 3). (B) Flow cytometry histograms comparing ITGα3 surface staining of parental cells, a CD151 KO clone with an ITGα3 surface export defect, and that CD151 KO clone transfected with a plasmid expressing wild-type CD151. (C) MFI plots comparing surface staining of CD151 KO clones defective in ITGα3 export to varying degrees, and those CD151 KO clones transfected with a plasmid expressing wild-type CD151. (B,C) Cells transfected with wild-type CD151 were gated for CD151 expression using the TS151 anti-CD151 antibody (Figure S7, related to Figure 3). (D) Effect of CD151 mutations on the ability of CD151 to rescue the ITGα3 surface export defect, assessed by flow cytometry. The extent of rescue is plotted as a function of mutation, based on the MFI of each sample (n ≥ 3). (E) Representative spinning disk confocal immunofluorescence images of CD151 KO clones showing variation in the degree of the ITGα3 surface export defect. (F) Immunofluorescence images of CD151 KO SVG-A cells transfected with CD151-mNeon, showing restoration of ITGα3 surface staining. (G) Adhesion of CD151 KO cells to plates coated with LmE8-332 after transfection with empty vector (ev), wild-type CD151, or the indicated CD151 mutant. Values are normalized to the maximum fraction of adherent (bound) cells in each experiment. Data are mean ± SEM of three independent experiments.

To further evaluate the basis for the ITGα3 surface staining defect in CD151 KO clones, we examined the localization of ITGα3 by immunofluorescence before and after transfection with wild-type CD151. Immunofluorescence staining revealed that ITGα3 is present in CD151 KOs but confined within intracellular puncta (Figure 3E), and that transfection of wild-type CD151 results in redistribution of ITGα3 to the cell surface and striking colocalization with CD151 (Figure 3F), indicating that CD151 promotes ITGα3 surface export.

### CD151 knockout leads to an ITGα3β1 adhesion defect

We adapted a cell-based assay to evaluate the effect of the CD151 knockout on integrin-dependent adhesion to a laminin-coated surface. The assay uses the E8 fragment of laminin-332 (LmE8), which retains the integrin-binding region of the protein and is sufficient to promote integrin-dependent cell adhesion (Figure S7E-G, related to Figure 3). Whereas parental SVG-A cells remain attached to LmE8-coated plates after washing (Figure 3G), TKO cells do not, indicating that cellular adhesion to the tissue culture surface is dependent on the presence of laminin-binding integrins. Likewise, CD151 knockout SVG-A cells with low ITGα3 surface staining also fail to stay attached to the plate (Figure 3G). The attachment defect in the CD151 knockout cells can be rescued by transfection of wild-type CD151 or by the non-perturbing Q202A mutant, but not by the R203E/H205E double mutant defective in integrin binding.

### CD151 and Tspan11 are functionally redundant

A number of different tetraspanins, such as CD63^12,13^, CD81^13,14^, and CD9^13,15^, have been implicated in binding to laminin-binding integrins in addition to CD151. To assess the potential of other tetraspanin proteins to bind ITGα3β1 and other laminin-binding integrins, we performed a computational screen using AlphaFold Multimer^16^ and the Structure Prediction and Omics informed Classifier (SPOC) Tool^17^ to identify high-confidence interactions (Figure 4A). This analysis accurately predicted the interaction of CD151 with ITGα3 at the membrane-proximal site seen in our experimental structure, as well as interactions of CD151 with ITGα6 and ITGα7 (Figure 4A and Figure S8, related to Figure 4), with high confidence (SPOC scores > 0.7). It also predicted with high confidence (SPOC scores >0.7) interactions of Tspan11 with ITGα6 and ITGα7, and moderate confidence (SPOC score >0.4) with ITGα3 at the same site, which is conserved between CD151 and Tspan11 (Figure 4A and Figure S8, related to Figure 4). An additional interaction between Tspan4 and ITGβ1 was also predicted with moderate confidence (Figure 4A).

**Figure 4.**
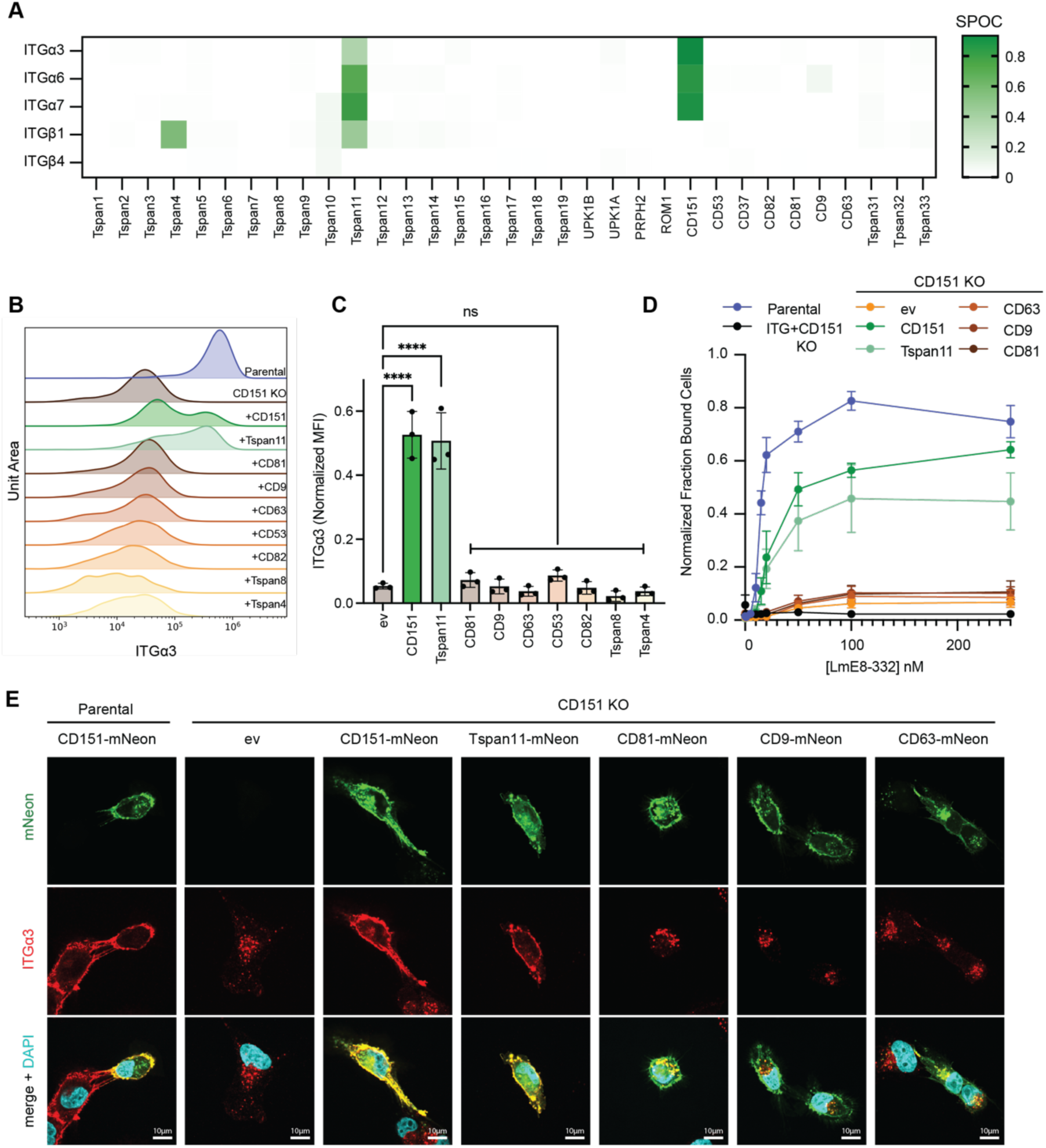
CD151 and Tspan11 rescue ITGα3 surface export and adhesion on laminin-332. (A) AlphaFold Multimer predictions of all potential complexes between laminin-binding integrin subunits and the 33 human tetraspanins were evaluated using the Structure Prediction and Omics-Classifier (SPOC), which ranks the likelihood of a true interaction on a scale from 0 (low probability) to 1 (high probability, green). (B) Analysis of the ability of different tetraspanin-mNeonGreen fusion proteins to rescue ITGα3 surface staining in CD151 KO SVG-A cells, as judged by flow cytometry. Histograms show the amount of ITGα3 staining after gating for mNeonGreen-positive cells (Figure S7, related to Figure 4). (C) Plot showing the mean fluorescent intensity (MFI) of ITGα3 surface staining, normalized to parental cells, for each tested tetraspanin-mNeonGreen fusion protein used to rescue CD151 KO cells. Each point on the plot represents the mean of four technical replicates (n = 3). Data was analyzed by one-way ANOVA, where ns, not significant, ****p<0.0001. (D) Adhesion of cells transfected with various tetraspanin-mNeonGreen fusion proteins to LmE8-332-coated plates. The fraction of cells that remained bound to the plate was quantified by measuring fluorescence at 510 nm of washed plates and normalization to the fluorescence at 510 nm of unwashed plates. Parental cells and TKO cells were transfected with CD151-mNeonGreen as reference samples. Data are plotted as the mean ± SEM of three independent experiments (n = 3). (E) Representative spinning disk confocal immunofluorescence images of parental or CD151 KO cells transfected with empty vector (ev) or with plasmids expressing various tetraspanin-mNeonGreen proteins.

We then made a panel of tetraspanins with mNeonGreen fused to the C-terminus to assess whether other tetraspanins could rescue ITGα3 surface export in CD151 knockout cells. We evaluated tetraspanins linked to integrin function (CD81, CD9, CD63, Tspan8, and CD82), predicted in the AlphaFold screen (Tspan4 and Tspan11) or not predicted to interact (CD53) as a negative control. Tspan11 rescued ITGα3 surface export as effectively as CD151, but all other tetraspanins failed to restore ITGα3 surface export, with activity not significantly different from cells transfected with empty vector (Figure 4B,C). Tspan11 also restored adhesion of CD151 knockout cells to LmE8-332 tissue culture plates (Figure 4D) and colocalized with ITGα3 at the plasma membrane (Figure 4E), whereas cells transfected with the other tetraspanins failed to restore adhesion on LmE8-332-coated plates (Figure 4D), and did not move ITGα3 out of intracellular puncta (Figure 4E).

## Discussion

In this study, we report the first example of a tetraspanin-integrin complex and establish a functional basis for the dependence of ITGα3β1 function on CD151. The interface between CD151 and ITGα3 is small (955 Å²) and dominated by the electrostatic complementarity of a basic patch on the tetraspanin and an acidic site on the calf-2 domain of ITGα3. The contact site is also distant from the ITGα3β1 headpiece and independent of ITGβ1, strongly suggesting that CD151 engagement will be unaffected by integrin conformation or by laminin binding, and arguing against a prior proposal that CD151 could conformationally modulate integrin affinity by promoting the high-affinity extended, open state.^18^ The location of the interaction site is fully consistent with a role for CD151 in integrin surface delivery, and contrasts with a recent claim that CD9 and CD81 can engage an integrin headpiece directly.^19^

Although prior studies did not report a reduction in surface ITGα3β1 after CD151 knockdown^18,20–25^, knockout^26^, or knockout in mice^27^, we did uncover a surface export defect in this work. A likely explanation for this discordance is that the surface defect was heterogeneous among individual knockout clones, and only by analyzing multiple individual clones in which CD151 was completely absent as a consequence of gene editing did the chaperone function of CD151 become clear.

Our mutagenesis data also provide strong support for a direct structure-function relationship dependent on the observed CD151-ITGα3 interface. The severity of the surface export and adhesion defects observed for individual CD151 point mutants tracked closely with the degree to which each mutation disrupted the electrostatic contacts identified in the structure: charge-reversal mutations at the core of the interface (K194E, H205E, and their combinations) were most disruptive, single alanine substitutions had intermediate effects, and mutations of residues outside the interface (Q202A, E213A, E213K) were functionally silent. This concordance among the biochemical co-immunoprecipitation results, cellular surface staining and adhesion data, and structural findings provides confidence that the interface we describe is the physiologically relevant determinant of CD151-dependent ITGα3β1 export. These results also suggest that the primary functional role of CD151 is to enable ITGα3β1 surface delivery, and that the that the adhesion defects we observe are a direct consequence of reduced receptor abundance.

Our findings have direct relevance for understanding the overlapping phenotypes of CD151 and ITGα3 loss of function, with loss of ITGα3β1 surface export providing a molecular mechanism for how CD151 silencing mutations also lead to epidermolysis bullosa phenotypes in humans (Figure 5). In the skin, ITGα3β1 and CD151 are expressed in the basal layer of the epidermis^4^ which binds laminin-332 in the basement membrane to preserve the structural integrity of the skin (Figure 5A). However, when CD151 is inactive, ITGα3β1 surface export is deficient, which impairs epidermal basal layer adhesion, leading to skin fragility and blistering (Figure 5B). Silencing mutations of ITGα3β1 have a severe phenotype in humans with skin fragility, renal abnormalities, and interstitial lung disease that often leads to death within a year, while missense mutations have less severe phenotypes with varying skin fragility and renal involement.^28^ Similarly, silencing of CD151 leads to localized skin fragility and neuropathy,^29^ and the variable penetrance of the export defect that we observed across independent CD151 knockout clones is consistent with the variable and often milder disease phenotype seen in patients with CD151 mutations relative to those with ITGα3 and laminin-332 loss of function.

**Figure 5.**
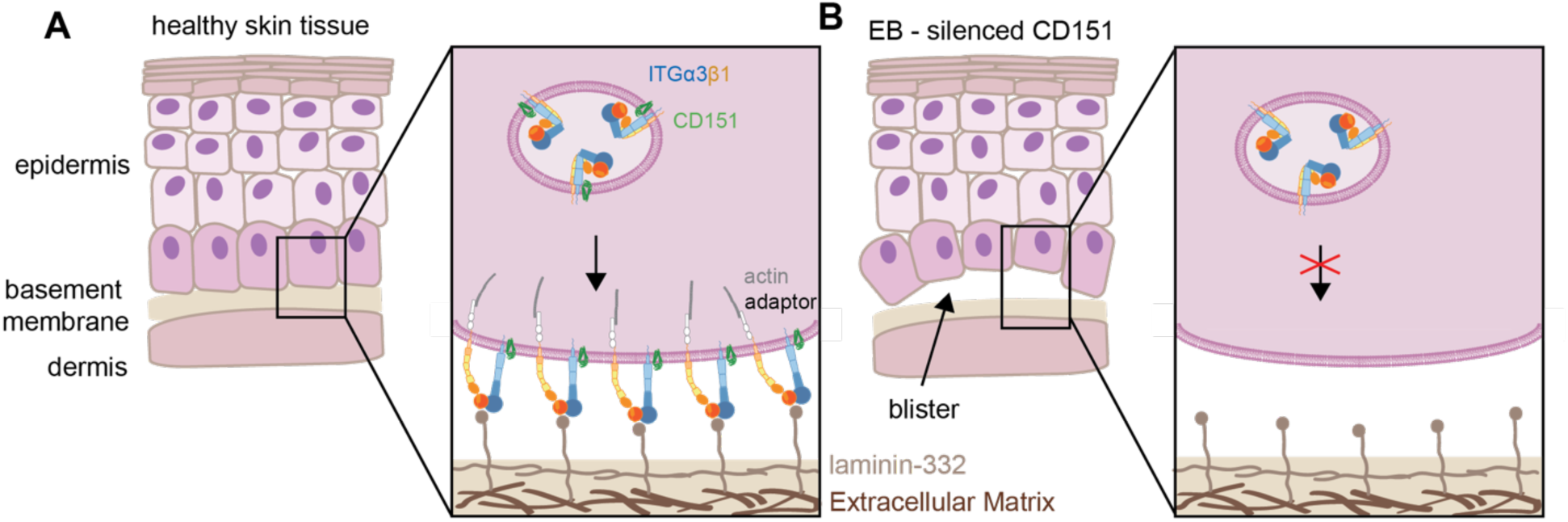
Proposed molecular mechanism of CD151-dependent epidermolysis bullosa. (A) Schematic of the layers of the skin from the dermis to the surface (left panel), showing attachment of the basal layer of the epidermis to the basement membrane in healthy tissue mediated by ITGα3β1 (right panel). (B) Schematic illustrating the effects of mutations in CD151 that lead to epidermolysis bullosa (EB). Loss of CD151 function results in blistering (right panel) because of impaired ITGα3β1 surface export (right panel) and defective adhesion of cells to the basement membrane.

One unanticipated finding of this study was that Tspan11 can compensate for the loss of CD151 in mediating ITGα3β1 surface delivery and adhesion to LmE8-coated plates. The residues at the ITGα3 contact interface with CD151 are conserved in Tspan11, but not in other tetraspanins reported to interact with laminin binding integrins.^3^ Tspan11 is a relatively understudied member of the tetraspanin family highly expressed in the colon and linked to focal adhesion assembly in osteoblasts.^30,31^ It is also possible that *TSPAN11* expression or genetic variation could act as a modifier of disease severity in patients with CD151-associated epidermolysis bullosa, as others have proposed.^32^

Lastly, recent cryo-EM structures of tetraspanin CD81 bound to the B cell co-receptor CD19 and of Tspan15 bound to the metalloprotease ADAM10 show that both ADAM10 and CD19 rely on interactions between the tetraspanin LEL and their membrane-proximal extracellular domains^5,33^ to facilitate surface delivery.^34,35^ Thus, the role of tetraspanins in chaperoning surface export of partner proteins through extracellular domain interactions appears to be an emerging broader principle in tetraspanin biology.

### Limitations of this study

The transmembrane and cytoplasmic regions of both ITGα3β1 and CD151 were not resolved in our reconstructions, leaving open the possibility that additional contacts within the membrane contribute to complex stability or to the effect of CD151 on integrin trafficking. While our data support a model in which CD151 acts primarily to promote surface export of ITGα3β1, we cannot exclude additional roles for CD151 in stabilizing or organizing the integrin after it arrives at the plasma membrane. In addition, the mechanistic basis for the variable penetrance of the export defect in independent CD151 knockout clones remains unexplained and could reflect clonal variability in Tspan11 or other compensatory factors, incomplete pathway dependence on CD151, or clonal differences unrelated to tetraspanin expression. Whether Tspan11 is responsible for the clonal variation in the export defect remains unresolved because the lack of availability of high-quality anti-Tspan11 antibodies makes detection and quantification of endogenous Tspan11 difficult. Future studies could be directed at defining the distribution of ITGα3β1 in the presence and absence of both CD151 and Tspan11, and at testing genetic interactions between CD151 and *TSPAN11 in vivo* to resolve this question. Finally, although the SPOC classifier improves predictive power beyond standard confidence metrics, AlphaFold Multimer remains a tool that still has potential for generating false positive (and false negative) interaction predictions.

## Methods

### ITGα3β1-CD151 Complex Expression and Purification

All constructs were cloned into pcDNA3.1-Zeo-TetO (tetR inducible) and expressed in Expi293T TetR inducible cells grown in Expi293 media (ThermoFisher). Constructs were transiently transfected with 1 µg DNA:0.8 µl FectoPro per ml culture volume (VWR). The fusion ITGα3-CD151 and ITGβ1 plasmids were transiently transfected at a ratio of 1:9 (w:w) respectively. Cells were transfected at 2.8×10^6^ cells/mL. 24 hrs after transfection, cells were enhanced with 3 mM valproic acid and 0.4% D-(+)-glucose (Sigma-Aldrich), and expression was induced with 0.004 mg/ml doxycycline. Cells were cultured for an additional 24 hrs, and then spun down at 4000 *g* for 15 min at 4 °C. Pellets were harvested, flash frozen, and stored at –80 °C.

Thawed pellets were lysed by hypotonic shock (20 mM HEPES pH 7.5, 2 mM MgCl_2_, 2 mg/mL iodoacetamide, 1:100,000 (v:v) benzonase, and EDTA free protease inhibitor), and mixed for 20 min at room temp. Cell membranes were harvested by centrifuging at 50,000 *g* for 15 min at 4 °C. Membranes were resuspended by dounce homogenization in solubilization buffer (20 mM HEPES pH 7.5, 250 mM NaCl, 1 mM CaCl_2_, 1% (w/v) lauryl maltose neopentyl glycol (LMNG), 0.1% (w/v) cholesteryl hemisuccinate tris salt (CHS)). The solubilization sample was mixed at 4 °C for 2-3 hours. Unsolubilized membranes were removed by centrifuging at 50,000 *g* for 45 min at 4 °C and passing the supernatant through a glass filter (VWR). The solubilized sample was passed over 4 ml M2 anti-Flag antibody resin at 4 °C.

The resin was washed with 50 mL Wash 1 buffer (20 mM HEPES pH 7.5, 250 mM NaCl, 1 mM CaCl_2_, 0.1% (w/v) LMNG, 0.01% (w/v) cholesteryl hemisuccinate), then slowly exchanged in 10% increments from Wash 1 to Wash 2 buffer (20 mM HEPES pH 7.5, 250 mM NaCl, 1 mM CaCl_2_, 0.05% (w/v) GDN, 0.005% (w/v) cholesteryl hemisuccinate). Sample was eluted with Wash 2 buffer + 0.2 mg/mL flag peptide. Elution fractions were concentrated using an Amicon concentrator with a 100 kDa or 50 kDa MW cutoff filter for full-length and truncated complexes, respectively. The concentrated sample was further purified by size exclusion chromatography (SEC) on a Sephadex S6i column. SEC peaks corresponding to the correct molecular weight were concentrated and evaluated by SDS-PAGE (Figure S1C, related to Figure 1).

The purified truncated complex was incubated with the TS151 anti-CD151 Fab^8^ and the anti-Fab nanobody^9^ at 5x molar excess for 1 hr at 4 °C. Fully formed complex with ITGα3-CD151, TS151 Fab and anti-Fab nanobody was purified by SEC and evaluated by SDS-PAGE (Figure S1D, related to Figure 1).

### TS151 Fab expression and purification

The heavy chain variable region of TS151 was cloned into the pFUSE-hIgG1-Fc2 vector (Invitrogen) with a 3C protease site inserted into the hinge region. The light chain variable region was cloned into the pD2610-v5 vector^34^. Expi293T cells were transiently transfected with both light chain and heavy chain vectors at a ratio of 1:3 (w:w) ratio. 24 hours after transfection cells were enhanced with 5 mM valproic acid and 0.4% D-(+)-glucose. After cell viability dropped below 50% (about 5-6 days) the media was harvested by centrifugation at 4000 *g* for 15 min at 4°C. Media was then loaded onto protein-A resin. The resin was washed with 50 mL HBS (20 mM HEPES pH 7.5, 150 mM NaCl). Bound antibody was eluted with 100 mM citrate pH 3.0, with immediately neutralization of the eluate in 1 M HEPES pH 8.0.

To generate the TS151 Fab, the purified TS151 antibody was incubated with 3C protease overnight at 4 °C. Complete cleavage was confirmed by SDS-PAGE. Free Fc was removed by passing the sample over the protein-A resin again, and the flowthrough was concentrated in a 30 kDa concentrator and buffer exchanged into Wash 2 buffer.

### Cryo-EM Grid Preparation

Immediately after complex formation, samples were concentrated and grids were prepared. The full-length complex was concentrated to 6.0 mg/mL and the truncated complex was concentrated to 5.5 mg/mL. Using a Vitrobot Mark IV set to 100% humidity at 8 °C, 4 µL of sample was applied to glow discharged UltrAuFoil 300 mesh grids, R 1.2/1.3 (TedPella). After a 10 s wait time, grids were blotted for 4 s with a blot force of 15 and then plunged in liquid ethane.

### Cryo-EM Data Collection

At the Harvard Cryo-Electron Microscopy Center for Structural Biology, grids were screened on a 200 kV Talos Artica (ThermoFisher), with a Volta phase plate and K3 detector (Gatan). Datasets were collected on a 300 kV Titan Krios G3i microscope, with a Falcon4i direct electron detector, Selectris Energy Filter, and Volta phase plate (ThermoFisher). Data were processed in CryoSparc, according to the workflow in Figures S2 and S3 (related to Figures 1,2).

For the full length ITGα3β1-CD151 complex, movies were motion corrected and dose weighted using Patch Motion Correction. Patch contrast transfer function (CTF) estimation was performed using Patch CTF Estimation. Particles were picked using blob picker and 2D classified. A subset of 2D classes was used for ab initio reconstruction model generation. Then all particles were used for multiple rounds of heterogeneous refinement. Particle orientation was rebalanced, reference-based motion correction was performed, and non-uniform refinement yielded the final full length map at a resolution of 3.84 Å. Because the headpiece was well resolved, we performed local refinement of the headpiece using a mask that included ITGα3 (33-610) and ITGβ1 (5-485) and yielded a final map at a resolution of 3.11 Å.

For the truncated CD151-ITGα3 complex, movies were motion corrected and dose weighted using Patch Motion Correction. Patch contrast transfer function (CTF) estimation was performed using Patch CTF Estimation. Particles were picked using blob picker and 2D classified. Two ab initio models were generated, and one contained all components of the complex. Particles were binned and eight rounds of heterogeneous refinement were performed. Particle orientation was rebalanced, binned particles were re-extracted, and local refinement yielded the final map at a resolution of 2.90 Å.

Atomic models of the ITGα3β1 headpiece and the CD151-ITGα3 calf-2 complex were built by docking AlphaFold 2 predicted models into the density map using ChimeraX followed by refinement using Phenix real-space refinement and Coot. In regions with backbone density but lacking sidechain density, protein sidechains are truncated to their C_β_.

### Cell Culture

SVG-A human fetal astrocyte cells were cultured at 37 °C and 5% CO_2_ in DMEM with 10% fetal bovine serum (FBS, GeminiBio) and 1% Penicillin-Streptomycin (PS, Thermo Fisher) and were confirmed to be negative for mycoplasma by PCR before use. For all assays, cells were detached from plates using 0.05% Trypsin/0.053 mM EDTA (Corning), a treatment confirmed not to proteolyze ITGα3.

### Genomic Engineering of SVG-A cells

Guide RNAs (gRNAs) targeting ITGα3, ITGα6, and CD151 were designed using CRISPOR. gRNAs targeting the N terminus with highest predicted efficiency and minimal off target effects were selected and cloned into the PX458 Cas9/GFP plasmid. SVG-A cells were transiently transfected using Lipofectamine 2000 (Invitrogen). Cells were washed 12 hours after transfection and fresh media was added. 2 days after transfection, cells were sorted using a Sony SH800 flow cytometer and cell sorter. GFP positive cells were single cell sorted into 96 well plates containing 40% conditioned DMEM, 40% fresh DMEM, and 20% FBS in each well. Single cell clones were expanded, and then each well was screened by western blot, flow cytometry, and/or genomic DNA PCR fragment sequencing to identify knockouts.

### CD151 Immunoprecipitation Assay

SVGA ITGα3, ITGα6, and CD151 triple KO clones were grown to ∼80% confluency in 6 well plates, and transiently transfected with 0.8 µg CD151, 1.6 µg ITGα3, and 0.8 µg empty pcDNA3.1 vector using 5 µL Lipofectamine 2000. 12 hours after transfection, cells were washed with fresh DMEM. Cells were harvested by centrifugation at 4000 *g* for 15 min. Pellets were washed with Phosphate Buffered Saline (PBS) (137 mM NaCl, 2.7 mM KCl, 10 mM Na_2_HPO_4_, 1.8 mM KPO_4_, pH 7.4) and then lysed by hypotonic shock in 750 µL of 20 mM HEPES buffer pH 7.5, containing 1 mM CaCl2, 2 mM MgCl2, 2 mg/mL iodoacetamide, 1:100,000 v:v benzonase. After mixing for 30 minutes at 4 °C, lysates were centrifuged at 21,000 *g* for 30 min at 4 °C to isolate membrane fractions. 50 µL RIPA buffer (150 mM NaCl, 50 mM Tris pH 8.0, 1% Triton, 0.5% sodium deoxycholate, 0.1% SDS, 1mM CaCl2) was then added to the membrane fractions and mixed for 2 hr at 4 °C, followed by centrifugation at 21,000 *g* for 45 min at 4 °C to remove unsolubilized material. The supernatant was applied to equilibrated anti-protein C affinity resin to bind the protein C affinity tag on ITGα3. The resin was mixed in an Eppendorf shaker at 26 °C, 500 rpm, for 1 hr. Resin was washed three times with 100 µL RIPA buffer and eluted in 20 µL RIPA buffer containing 0.2 mg/mL protein C peptide and 5 mM EDTA. The amounts of CD151 in the input and immunoprecipitated samples were quantified by western blot, using an α-CD151 TS151 antibody. Band intensity was quantified in Image Lab (BioRad) and normalized to wild type CD151.

### Cell Surface Export Assay

7.5 × 10^4^ CD151 KO SVG-A cells were seeded at roughly 70% confluency in each well of a 24 well plate and transfected 24 hours later. For analysis of wild-type and mutant CD151-facilitated export, the transfection mix included 1 µL Lipofectamine 2000 (Invitrogen), 0.5 µg of empty pcDNA3 vector and 0.5 µg of a pcDNA3 plasmid expressing wild-type CD151 or mutant CD151. Cells were washed after 12 hr, trypsinized and harvested at 48 hr, and then transferred into individual wells of a 96 well plate at 1.0 × 10^5^ cells/well. Cells were washed with PBS, then stained with anti-CD151 TS151-Alexa Fluor 488 at 1:400 dilution in HBS containing 0.1% BSA to identify transfected cells and with anti-ITGα3-APC antibody (Biolegend) at 1:400 in HBS containing 0.1% bovine serum albumin (BSA, Roche) to quantify export. Samples were analyzed on a CytoFLEX flow cytometer (Beckman Coulter) gating for positive fluorescence in the anti-TS151 channel (Figure S7, related to Figures 3,4). For analysis of other tetraspanin proteins, a family of mNeonGreen fusion proteins were constructed, transfections were carried out using 1 µL Lipofectamine 2000 (Invitrogen), 0.5 µg of empty pcDNA3 vector, and 0.5 µg of the tetraspanin-mNeonGreen fusion-expressing plasmid, and cells were gated for positive tetraspanin expression by mNeonGreen fluorescence. Analysis of all samples was carried out using EasyFlow Q software^36^, normalized to SVG-A ITGα3 surface staining in parental cells.

### Laminin E8 Fragment Design, Expression, and Purification

Laminin-332 E8 fragment (LmE8-332) containing the minimal integrin binding domain was designed by aligning to the sequences of Lm-E8-111 and Lm-E8-511 complexes for which structures had been reported.^37,38^ Sequences aligned well in the coiled-coiled domain, and LmE8-332 constructs were designed to match. Cysteine mutations were introduced into the coiled-coiled domain at sites that aligned well to those introduced in Lm-E8-511.^37^ Subunits were cloned into pTarget with N-terminal HA secretion sequences and FLAG, HA, and His_6_ tags for alpha, beta, and gamma subunits, respectively. Constructs were transiently transfected at a ratio of 1:1:1 (w:w:w) in Expi293T cells. 24 hours after transfection cells were enhanced with 5 mM valproic acid and 0.4% D-(+)-glucose. After cell viability dropped below 50% (about 5-6 days) the media was harvested by centrifugation at 4000 *g* for 15 min at 4 °C. The resulting supernatant was supplemented with 1 mM CaCl_2_ and run over M1-Flag affinity resin. Resin was washed twice with 50 mL HBS containing 1 mM CaCl_2_, and protein was eluted with HBS containing 0.1 mg/mL flag peptide and 5 mM EDTA. The eluate was concentrated using an Amicon concentrator with a 50 kDa cutoff filter and then purified by SEC on a S6i column. The SEC peak contained all 3 subunits as judged by SDS-PAGE and western blot.

### Adhesion Assay using LmE8-332

SVG-A cells were seeded in 24 well format at roughly 70% confluency and were transfected after 24 hr incubation. For analysis of cellular adhesion in cells expressing wild-type or mutant CD151, the transfection mix included 1 µL Lipofectamine 2000 (Invitrogen), 0.5 µg of a pcDNA3 plasmid expressing wild-type CD151 or mutant CD151, and 0.5 µg of a pcDNA3 plasmid expressing GFP to identify transfected cells. For analysis of other tetraspanin proteins, a family of mNeonGreen fusion proteins were constructed, and transfections were carried out using 1 µL Lipofectamine 2000 (Invitrogen), 0.5 µg of empty pcDNA3 vector, and 0.5 µg of the tetraspanin-mNeonGreen fusion-expressing plasmid. Laminin was then immobilized on 96 well high binding black/clear bottom plates (Corning) by adding 100µL LmE8-332 solution and incubating at 4 °C overnight. The next day, plates were washed twice with PBS and blocked with PBS containing 30 mg/mL BSA (Roche) at 37 °C for 1 hr. Then plates were washed twice with PBS. SVG-A cells were trypsinized from the 24 well plates for 5 min and diluted to 2.0 × 10^5^ cells/mL in DMEM lacking phenol red (ThermoFisher). 100 µL of diluted cells were added to each well and allowed to adhere for 1 hr at 37 °C, 5% CO_2_. Cells were washed 4 times using a BioTEK plate washer and resuspended in PBS. GFP fluorescence was measured on a SpectraMax analyzer using 488 nm excitation and 515 nm emission. Background signal was subtracted and the fraction of cells bound was calculated from the change in GFP fluorescence as a result of washing, using an unwashed plate as a reference. Data were normalized by setting the maximum fraction bound cells to 1.

### Immunofluorescence Imaging

Washed cover slides (Warner Instruments) were prepared with 10 µg/mL poly-D-lysine diluted in PBS and incubated overnight at 4 °C in a 6 well plate. Slides were washed three times with PBS, and 1.0 × 10^5^ cells in 2 mL DMEM were added to each slide. Cells were incubated on these slides overnight. The next day, slides were washed 3 times with PBS, fixed with 4% paraformaldehyde in PBS for 15 min at room temperature (RT), washed 3 times with PBS, quenched with 0.1 M glycine in water for 15 min at RT, washed 3 times with PBS, permeabilized with 0.1% Triton X-100 in PBS for 10 min at RT, washed 3 times with PBS, and blocked with 3% BSA (Roche) in PBS for 1 hr at RT in the dark. α-ITGα3 antibody (Biolegend) was diluted to 0.001 mg/mL in blocking buffer and added to slides for 1 hr at RT in the dark. Slides were washed 2 times with PBS, then DAPI diluted 1:1000 (Sigma-Aldrich) was added to slides for 5 min at RT in the dark and slides were washed 3 times with PBS. Slides were mounted using Prolong Diamond Antifade Mountant (Thermo Fisher), which was left to dry overnight at RT in the dark. Slides were then stored at 4 °C in the dark. Slides were imaged at the Harvard Core for Imaging Technology and Education on a spinning disk confocal microscope, and images were analyzed using Image J (FIJI).

## Acknowledgments

We thank the cryo-EM Center for Structural Biology at Harvard Medical School for expert assistance with cryo-EM data acquisition and processing. We thank the Core for Imaging Technology and Education for expert assistance with cell imaging acquisition. We also thank Yuxin Hao and the Springer lab for advice on cell adhesion assay protocols. We thank Tim Springer for advice on integrin truncation approaches. And we thank Katherine Susa for advice on project conception and shared tetraspanin expertise. Financial support for this work was provided by NIH grant R01 AI172846 (to S.C.B.), a gift from Edward B. Goodnow (to S.C.B.), and F31 AR083740 to LL. We also thank members of the Blacklow and Kruse labs for helpful discussions.

## Author contributions

S.C.B. A.C.K., and L.L conceived of the research. S.C.B. A.C.K., and L.L designed the experiments. L.L. performed experiments. E.W. prepared CD151 knockout clones. S.C.B., A.C.K., and L.L interpreted the data. S.C.B. and L.L wrote the manuscript with input from A.C.K. in the editing phase.

## Declaration of interests

S.C.B. is on the board of directors for the non-profit Revson Foundation and non-profit Institute for Protein Innovation, is on the scientific advisory board for Erasca, Inc., and is head of the scientific advisory board for Odyssey Therapeutics. A.C.K. is a co-founder and consultant for Tectonic Therapeutic and Seismic Therapeutic and for the Institute for Protein Innovation, a non-profit research institute.

---

Key resources table.

Key resources table
| REAGENT or RESOURCE | SOURCE | IDENTIFIER |
| --- | --- | --- |
| **Antibodies** |  |  |
| APC anti-ITGα3 | Biologend | Cat# 343808 |
| anti-CD151 (TS151) | Produced in-house | N/A |
| Alexa Fluor 488 anti-CD151 (TS151) | Produced in-house | N/A |
| anti-CD151 | Cell Signaling Technology | Cat# 96282S |
| anti-β tubulin | Cell Signaling Technology | Cat# 86298S |
| APC anti-ITGα6 | Biologend | Cat# 313615 |
| **Bacterial and virus strains** |  |  |
| DH5α | Thermo Fisher Scientific | Cat# 18258-012 |
| **Biological samples** |  |  |
| **Chemicals, peptides, and recombinant proteins** |  |  |
| FBS | GeminiBio | Cat# 100-106 |
| Penicillin and streptomycin | Thermo Fisher Scientific | Cat# 15140163 |
| Trypsin/0.53 mM EDTA in HBSS | Corning | Cat# 25-051-CI |
| DMEM | Corning | Cat# 10-017-CV |
| DMEM, no phenol red | Thermo Fisher Scientific | Cat# 21-063-029 |
| Expi293 expression medium | Thermo Fisher Scientific | Cat # A1435103 |
| Opti-MEM | Life Technologies | Cat# 31985-070 |
| Doxycycline Hyclate | Sigma-Aldrich | Cat# D9891 |
| PBS | Corning | Cat# 21-031-CV |
| NaCl | VWR | Cat# 0241-10KG |
| HEPES | Sigma-Aldrich | Cat# H4034-1KG |
| Sodium deoxycholate | Sigma-Aldrich | Cat# D6750 |
| SDS | Sigma | Cat # 75746 |
| lauryl maltose neopentyl glycol | Anatrace | Cat# NG310 |
| GDN | Anatrace | Cat# GDN10 |
| cholesteryl hemisuccinate tris salt | Anatrace | Cat# 102601-49-0 |
| EDTA free protease inhibitor | Thermo Fisher Scientific | Cat# A32965 |
| EDTA | Sigma-Aldrich | Cat# E9884 |
| Iodoacetimide | Sigma-Aldrich | Cat# I1149 |
| Flag peptide (DYKDDDK) | GenScript | N/A |
| Protein-C peptide (EDQVDPRLIDGK) | GenScript | N/A |
| Triton X-100 | Sigma-Aldrich | Cat# T8787-250ML |
| 16% paraformaldehyde | Thermo Fisher Scientific | Cat# 047340-9M |
| Bovine Serum Albumin | Sigma-Aldrich | Cat# 10735094001 |
| Poly-D-Lysine | Thermo Fisher Scientific | Cat# A3890401 |
| Trizma base | Sigma-Aldrich | Cat# T1503 |
| Glycine | Sigma-Aldrich | Cat# G7126 |
| Valproic acid sodium salt | Sigma-Aldrich | Cat# P4543 |
| D-(+)-Glucose solution | Sigma-Aldrich | Cat# G8769 |
| Alexa Fluor 488 NHS ester | Thermo Fisher Scientific | Cat# A20000 |
| DAPI | Sigma-Aldrich | Cat# MBD0015 |
| M2 anti-Flag resin | Produced in-house | N/A |
| M1 anti-Flag resin | Produced in-house | N/A |
| anti-protein C resin | Produced in-house | N/A |
| 3C protease | Produced in-house | N/A |
| **Critical commercial assays** |  |  |
| QIAprep Spin Miniprep Kit | Qiagen | Cat# 27106 |
| ZymoPURE Maxiprep Kit | Zymo Research | Cat# D4203 |
| KOD1 PCR Master Mix | Milipore Sigma | Cat# KMM-201NV |
| NEBuilder HiFi DNA Assembly | New England Biolabs | Cat# E2621L |
| QuickChange Lightning Mutagenesis Kit | Aligent | Cat# 210513 |
| Benzonase | Millipore Sigma | Cat# E1014-25KU |
| Polyplus-transfection FectoPRO® | VWR | Cat# 10118-842 |
| Lipofectamine™ 2000 | Thermo Fisher Scientific | Cat# 11668019 |
| Superose 6 Increase 10/300 GL | Cytiva | Cat# 29091596 |
| Prolong Diamond Antifade Mountant | Thermo Fisher Scientific | Cat# P36961 |
| **Deposited data** |  |  |
| EM density map of the full-length CD151-ITGα3β1 complex | This paper | EMDB: EMD-78902 |
| EM density map and atomic coordinates for the ITGα3β1 complex | This paper | EMDB: EMD-78903<br>PDB: 38LD |
| EM density map and atomic coordinates for the CD151-ITGα3 calf-2 truncated protein | This paper | EMDB: EMD-78901<br>PDB: 38LC |
| **Experimental models: Cell lines** |  |  |
| Human: Expi293F Inducible cells | Thermo Fisher Scientific | Cat# A39241 |
| Human: SVG-A cells | Walter J. Atwood, Brown University | RRID: CVCL_5G13 |
| SVGA CD151 KO cells | This paper | N/A |
| SVGA ITGα3, ITGα6, and CD151 TKO cells | This paper | N/A |
| **Oligonucleotides: oligonucleotides for this study are listed in Table S1.** |  |  |
| **Recombinant DNA** |  |  |
| pCDNA3.1(+) | Thermo Fisher Scientific | Cat# V79020 |
| pCDNA3.1(+)-Zeo-tetO | Thermo Fisher Scientific | Cat# V86020 |
| pFUSE-hlgG1-Fc | Invivogen | Cat code: pfuse-hg1fc1 |
| pD2610-v5 | ATUM | N/A |
| pX458 | Addgene, pSpCas9(BB)-2A-GFP (PX458) was a gift from Feng Zhang | RRID: Addgene_48138 |
| pTarget | Addgene | RRID:<br>Addgene_186452 |
| mNeonGreen | Addgene, NeonGreen-Giantin | RRID:<br>Addgene_98880 |
| CD151 | Addgene | RRID:<br>Addgene_58079 |
| ITGa3 | DNASU | RRID:<br>HsCD00859796 |
| ITGβ1 | Addgene | RRID:<br>Addgene_ |
| GFP | Addgene | RRID:<br>Addgene_11150 |
| Synthesized DNA fragments | Twist Bioscience | N/A |
| **Software and algorithms** |  |  |
| CryoSparc v3, v4 | Punjani et al. | https://www.cryosparc.com |
| Phenix | Adams et al. | https://sbgrid.org/software/ |
| Coot | Emsley and Cowtan | https://sbgrid.org/software/ |
| UCSF ChimeraX 1.5 | Pettersen et al. | https://www.cgl.ucsf.edu/chimerax/ |
| PDB validation server | World Wide Protein Data Bank | https://www.wwpdb.org/ |
| GraphPad Prism | GraphPad | RRID:SCR_002798 |
| ImageLab | Bio Rad | https://www.bio-rad.com/ |
| EasyFlowQ | Ma et al. | https://ym3141.github.io/EasyFlowQ/ |
| FIJI | Schindelin et al. | https://imagej.net/Fiji ; RRID: SCR_002285 |
| **Other** |  |  |
| UltraAuFoil R 1.2/1.3, 300 mesh, gold | Electron Microscopy Sciences | Cat# Q350AR13A |
| 25 mm cover slips | Warner Instruments | Cat# CS-25R17 |
| Poly-Prep Slides | Milipore Sigma | Cat# P0425 |
| Protran nitrocellulose membrane | Cytiva | Cat# 10600010 |
| Amicon Centrifugal filter, 50 kDa | Millipore Sigma | Cat# UFC905024 |
| Amicon Centrifugal filter, 100 kDa | Millipore Sigma | Cat# UFC910024 |

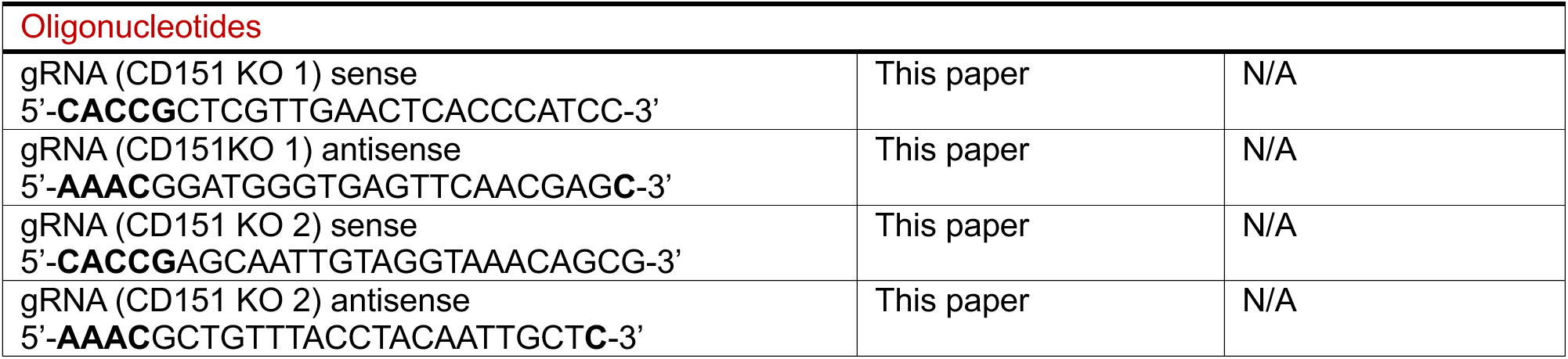
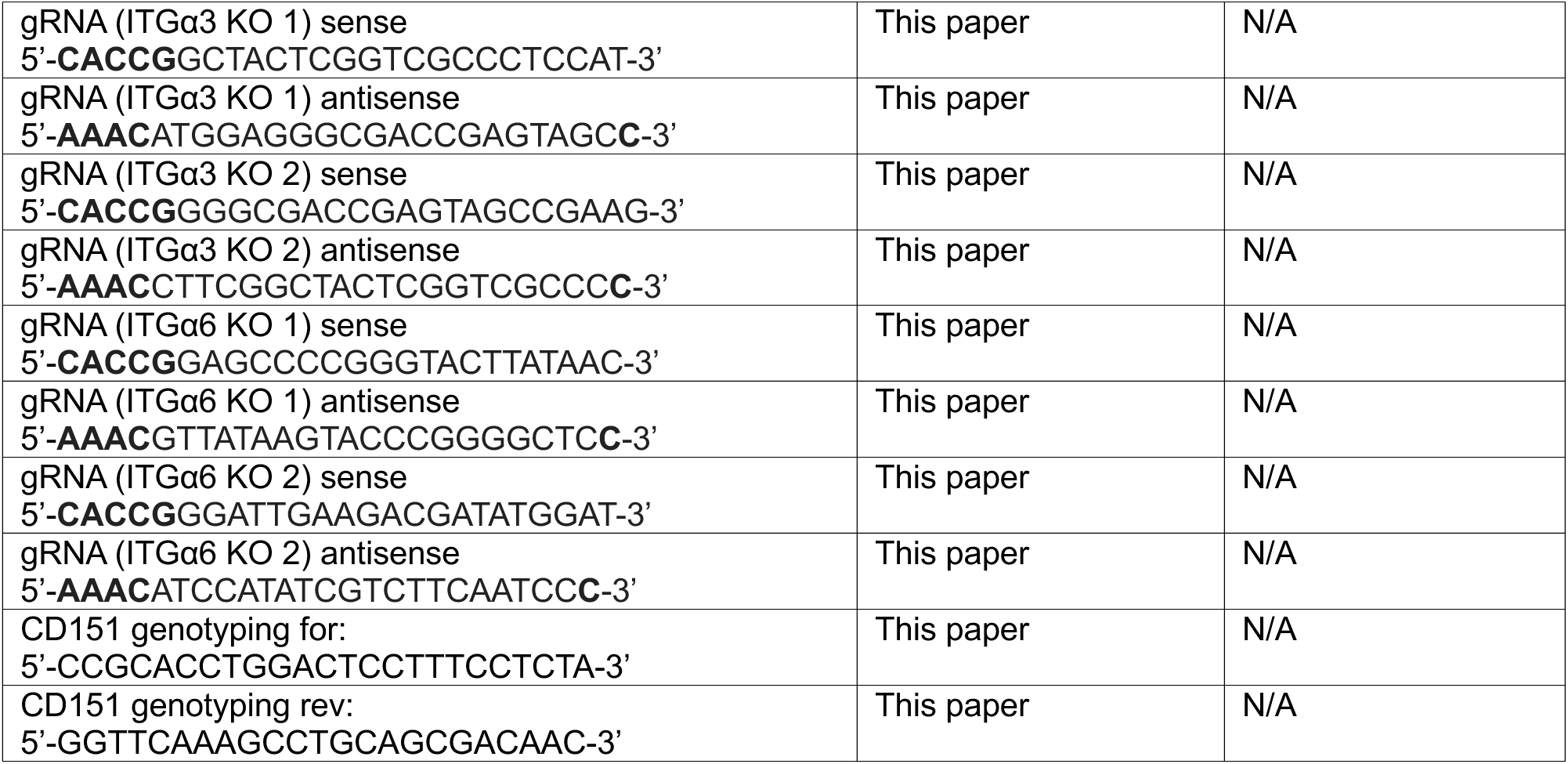
Table S1.

**Table S2.**
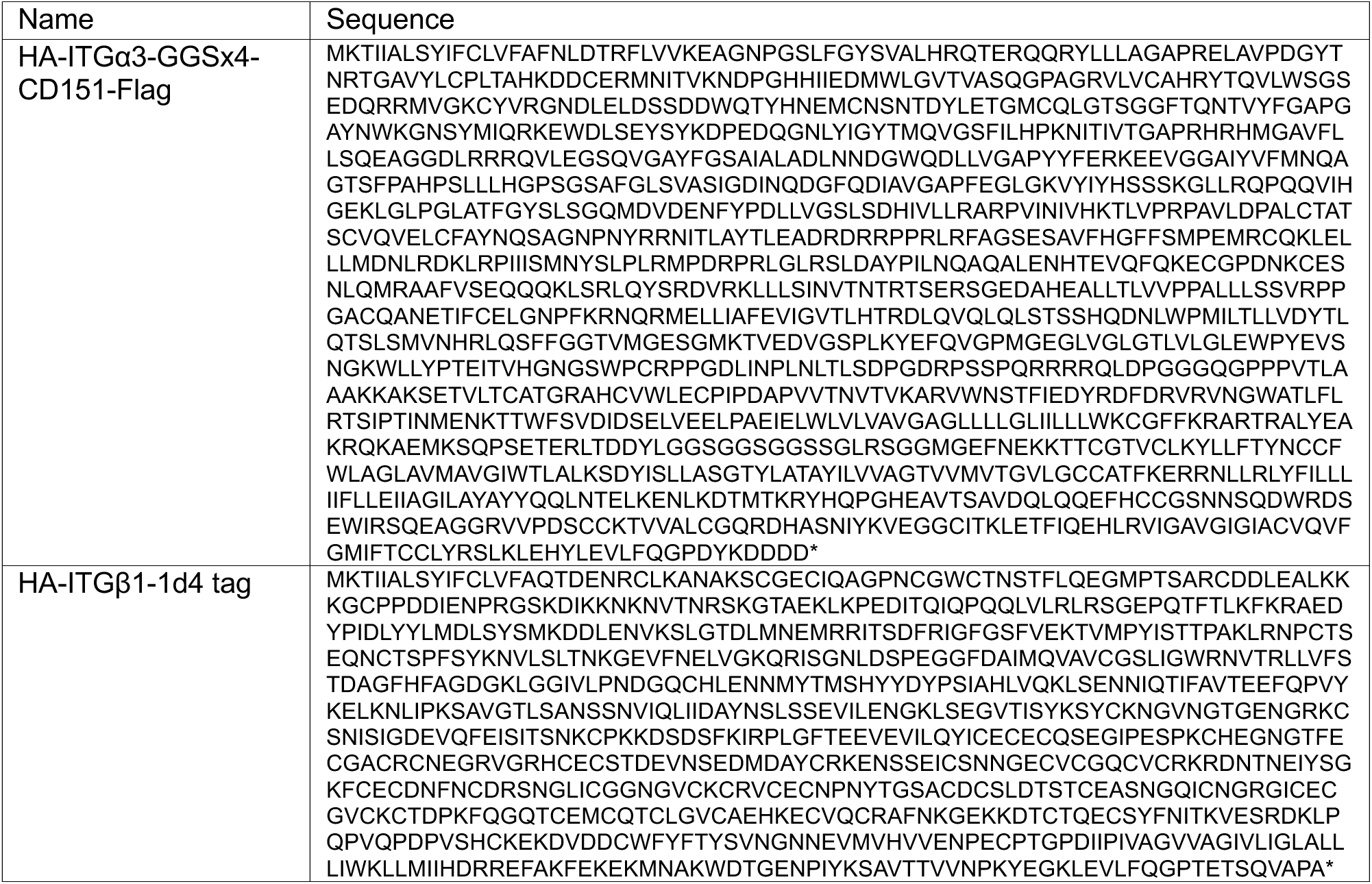
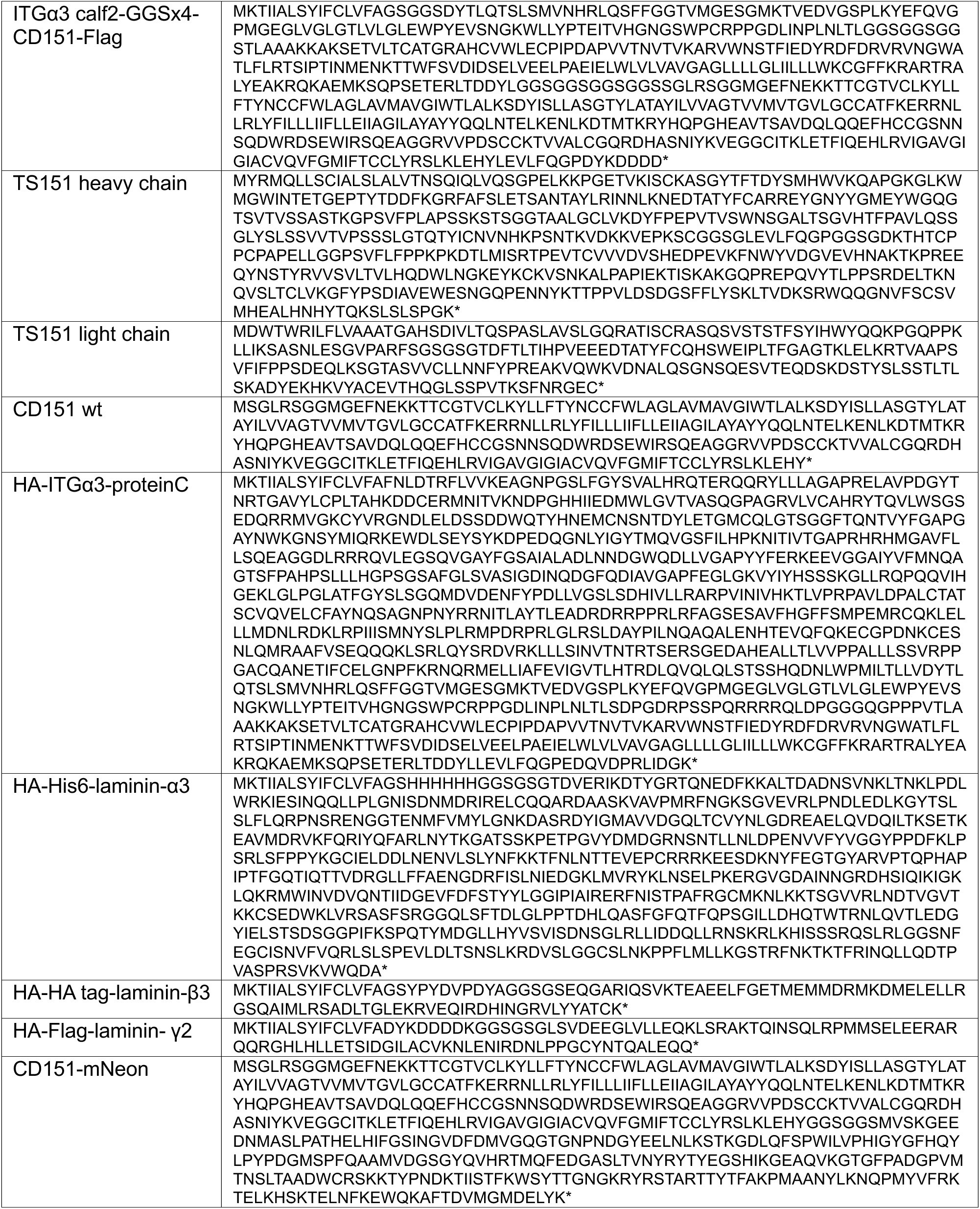
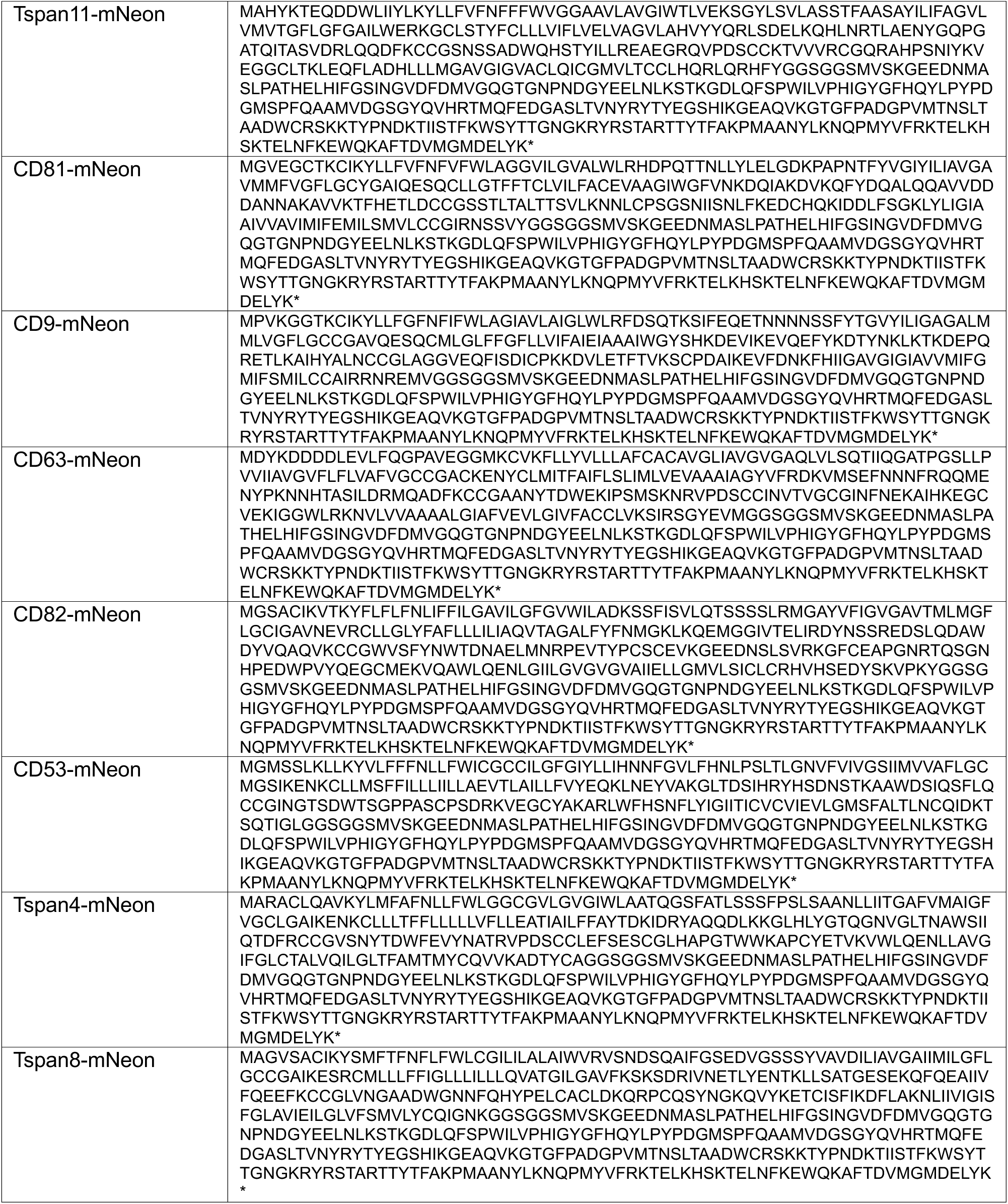
Protein Sequences.

**Figure S1.**
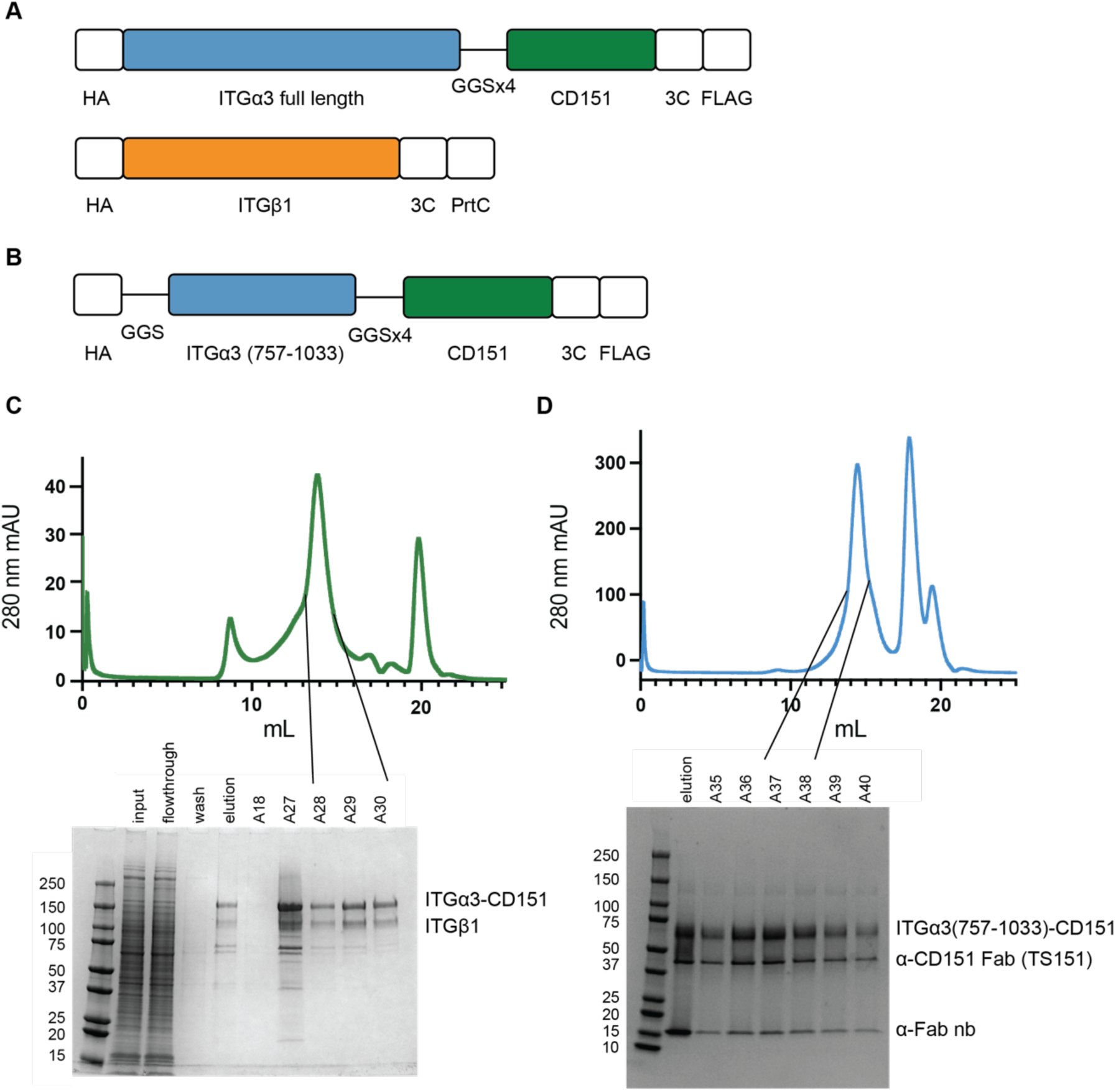
CD151-ITGα3β1 construct design and purification. (A, B) Schematics showing proteins used for protein expression and complex formation. HA: hemagglutinin signal sequence; 3C: 3C protease cleavage site; FLAG, FLAG affinity tag; PrtC, protein-C affinity tag. The ITGα3 truncated construct has an additional GGS linker after the HA signal sequence. (C) Size exclusion chromatogram of the CD151-ITGα3β1 full length complex after affinity chromatography, accompanied by a Coomassie-stained gel showing sample purity in fractions A28-A30 used for cryoEM sample preparation. (D) Size exclusion chromatogram of the CD151-ITGα3 calf-2 complex bound to the TS151 antibody and α-Fab nanobody after affinity chromatography, accompanied by a Coomassie-stained gel showing sample purity in fractions A37-A38 used for cryoEM sample preparation.

**Figure S2.**
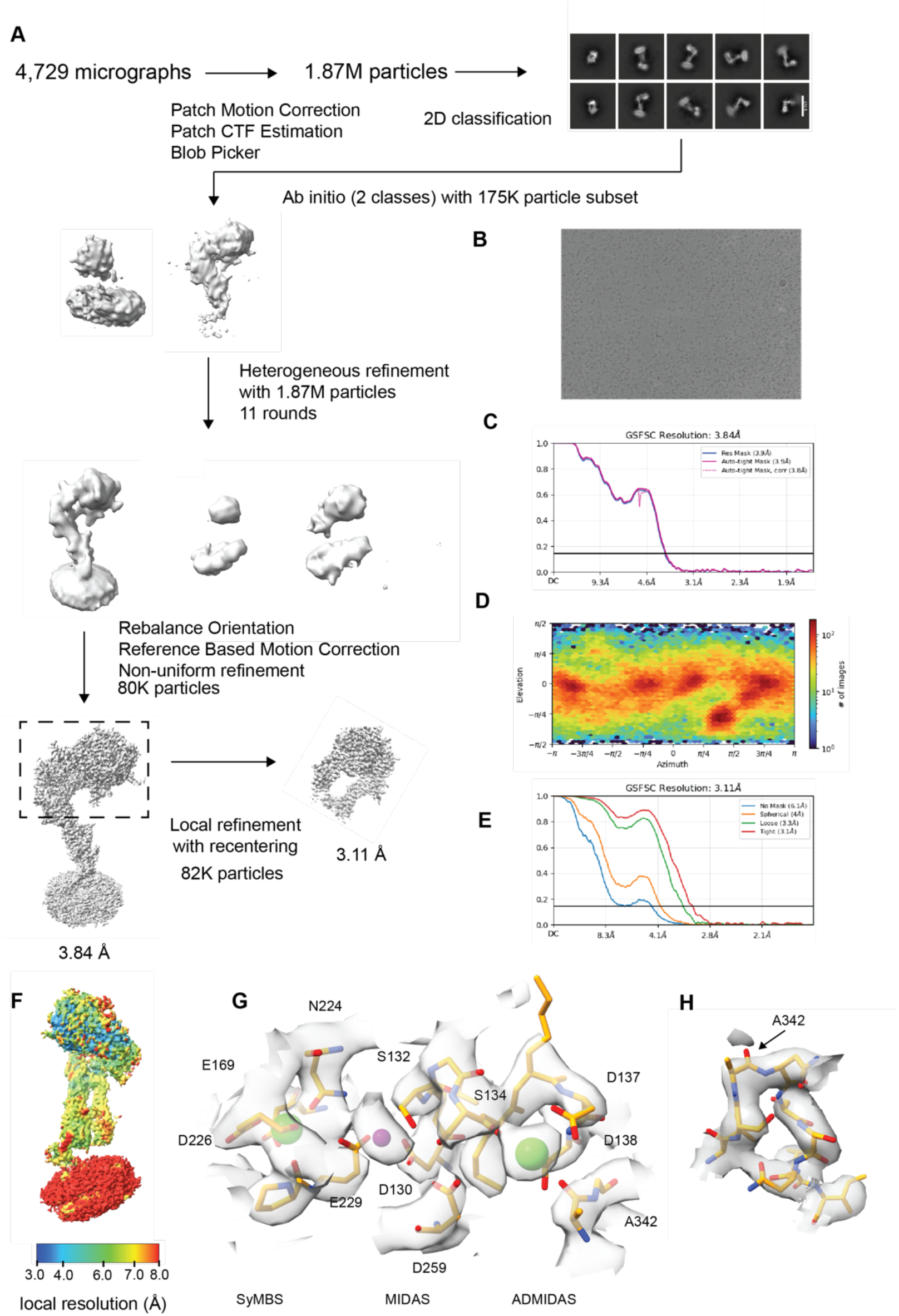
Cryo-EM data and processing workflow for the full length CD151-ITGα3β1 complex. (A) Image processing workflow. Milestone maps are shown to display progression in map quality during processing. (B) Representative micrograph image. (C) Fourier shell correlation (FSC) curve used to estimate resolution of the full CD151-ITGα3β1 reconstruction from the map. (D) Orientation distribution of particles used in final refinements. (E) Fourier shell correlation (FSC) curve used to estimate resolution of the ITGα3β1 headpiece reconstruction from the map. (F) Local resolution of CD151-ITGα3β1 and ITGα3β1 headpiece maps, colored on a sliding scale from blue (highest resolution) to red (lowest resolution). (G) Cryo-EM density map and model for ITGβ1 MIDAS, ADMIDAS, and SyMBS sites. (H) Cryo-EM density map and model for the α6-β7 loop of ITGβ1.

**Figure S3.**
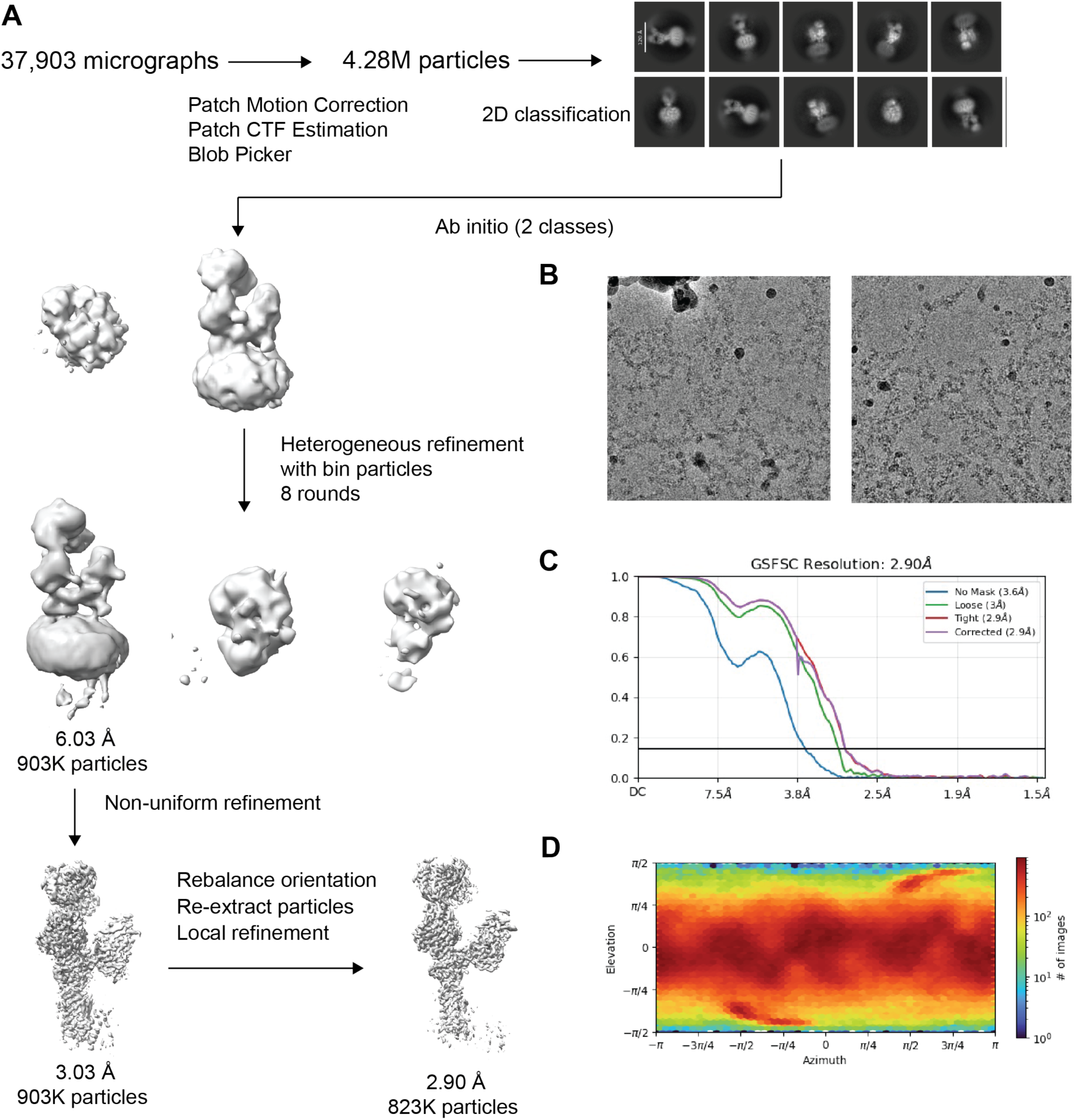
Cryo-EM data processing of CD151-ITGα3 calf-2 complex. (A) Image processing workflow. Milestone maps are shown to display progression in map quality during processing. (B) Representative micrograph images. (C) Fourier shell correlation (FSC) curve used to estimate resolution of the CD151-ITGα3 calf-2 reconstruction from the map. (D) Orientation distribution of particles used in final refinements.

**Figure S4.**
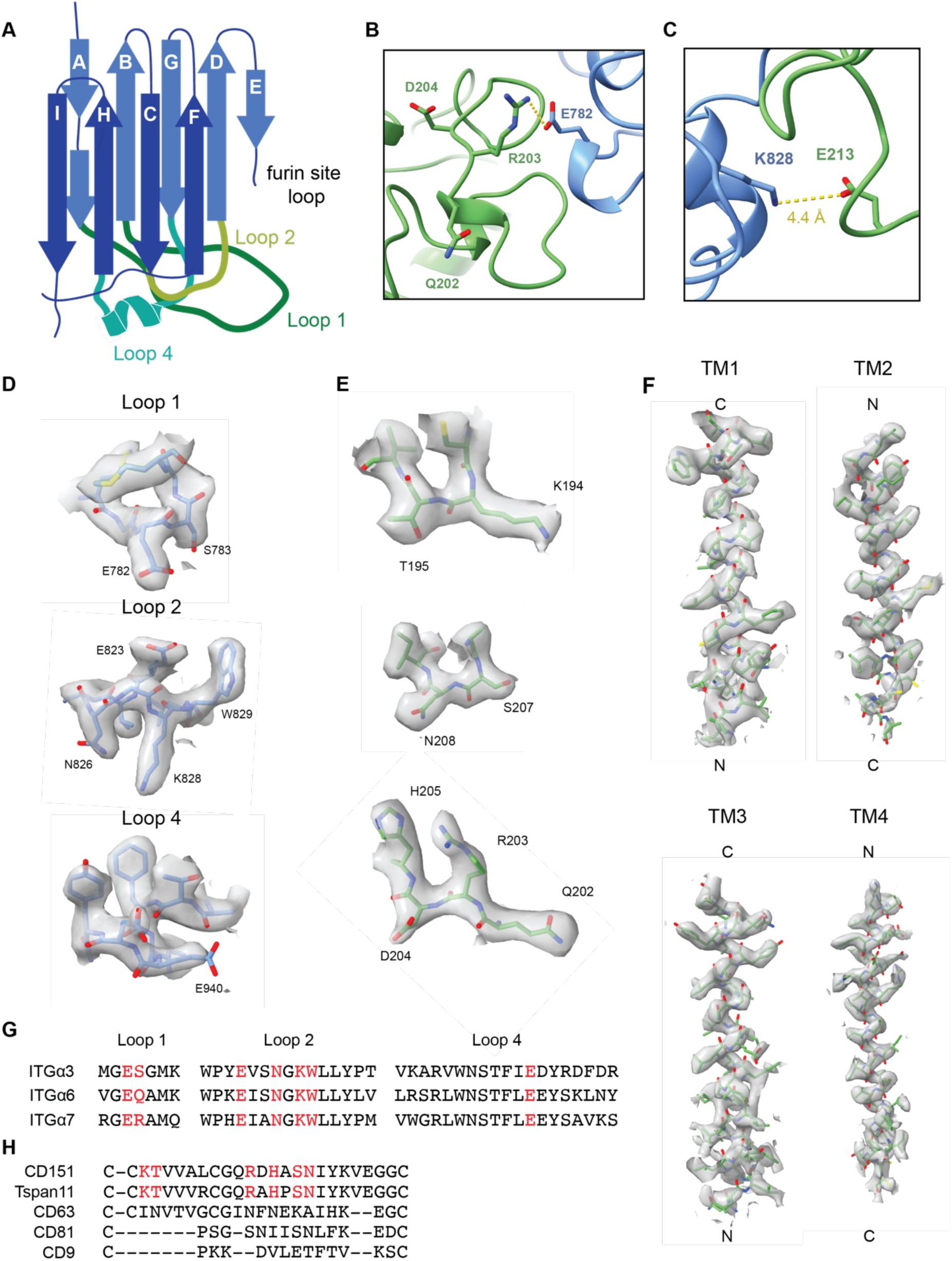
CD151-ITGα3 calf-2 modeling. (A) Topology diagram of ITGα3 calf-2 (blue), showing secondary structure with CD151-interacting loops highlighted in various shades of green. (B) CD151-ITGα3 interface highlighting the CD151 “QRD” motif (sidechains rendered as sticks), indicating that only R203 makes direct contact with ITGα3 residue E782. CD151 is green, and ITGα3 is blue. (C) CD151-ITGα3 interface highlighting the side chain contact between N208 of CD151 with S783 of ITGα3. CD151 is green, and ITGα3 is blue. (D-F) Cryo-EM density and accompanying atomic model for ITGα3 calf-2 loops (D), CD151 extracellular segments in contact with ITGα3 (E), and the transmembrane domain of CD151 (F). (G) Sequence alignment of laminin-binding integrin calf-2 loops, highlighting ITGα3 residues that contact CD151, and the analogous positions in ITGα6 and ITGα7, in red. (H) Sequence alignment of selected tetraspanins in the region of CD151 that contacts ITGα3. Residues that contact ITGα3, and the analogous positions of the aligned tetraspanins, are highlighted in red.

**Figure S5.**
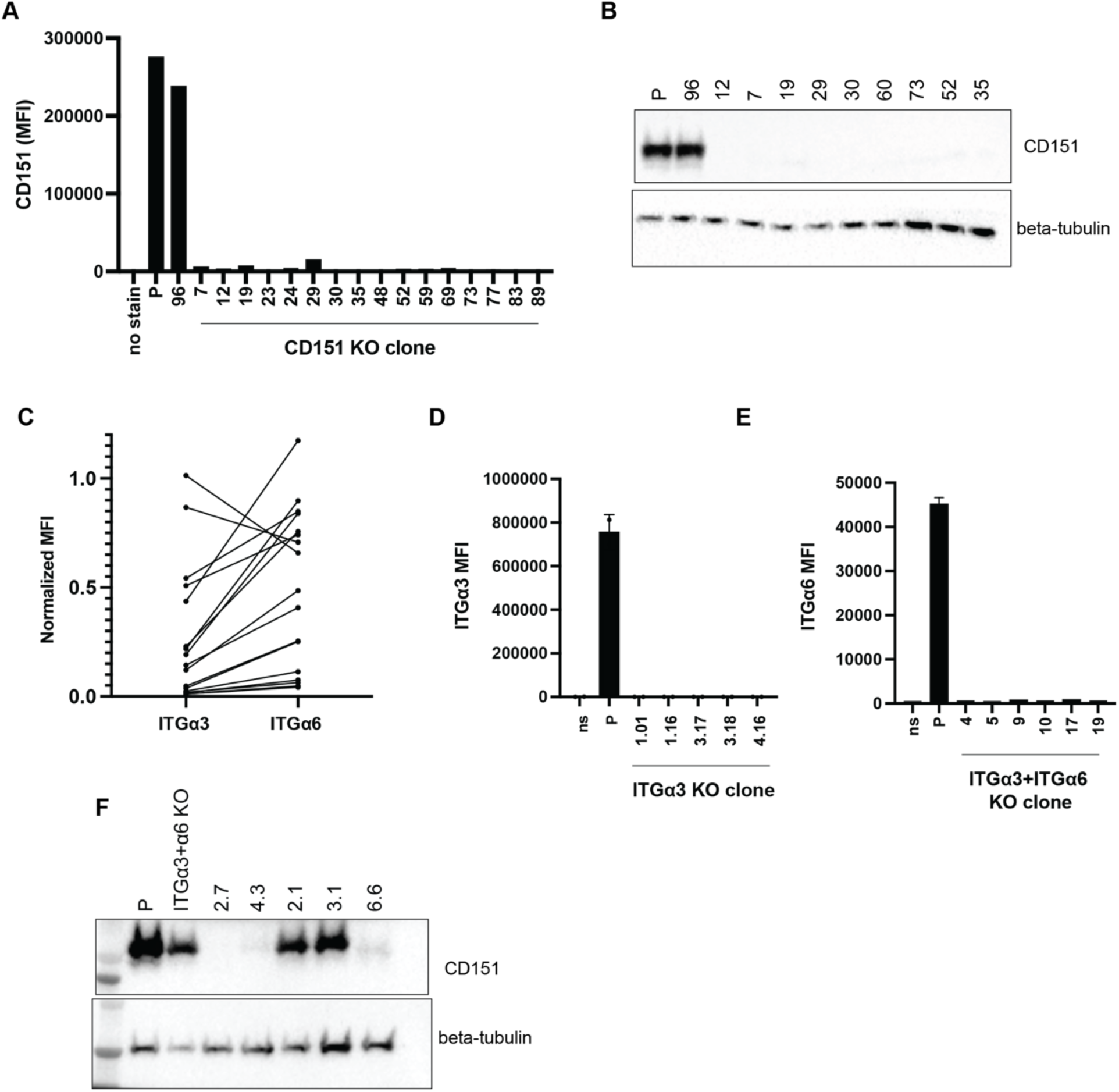
Identification of SVG-A KO clones. (A) Candidate CD151 KO clones were expanded and then screened by flow cytometry to identify knockouts. Clone 96, which did not diminish CD151 surface staining, was retained as a CRISPR/Cas9-treated parental control. (B) Western blot analysis to confirm CD151 knockout in candidate clones. Genomic PCR and sequencing confirmed KO phenotypes with various frameshifts and deletions. (C) Correlation of surface staining (MFI, normalized to parental control) reductions for ITGα3 and ITGα6 in CD151 KO clones, as judged by flow cytometry, showing that loss of surface ITGα3 was highly correlated with loss of surface ITGα6 in clones with the strongest export defects. (D) Candidate ITGα3 KO clones were expanded and then screened by flow cytometry to identify knockouts. (E) ITGα3 KO SVG-A cells were treated with an ITGα6 CRISPR/Cas9 targeting vector to create candidate ITGα3/ITGα6 DKO clones. The candidate DKO clones were expanded and then screened by flow cytometry to identify DKO lines. (F) ITGα3/ITGα6 DKO clones were treated with an CD151 CRISPR/Cas9 targeting vector to create candidate TKO clones. CD151 knockout in the TKO lines was confirmed by Western blot, using the TS151 antibody.

**Figure S6.**
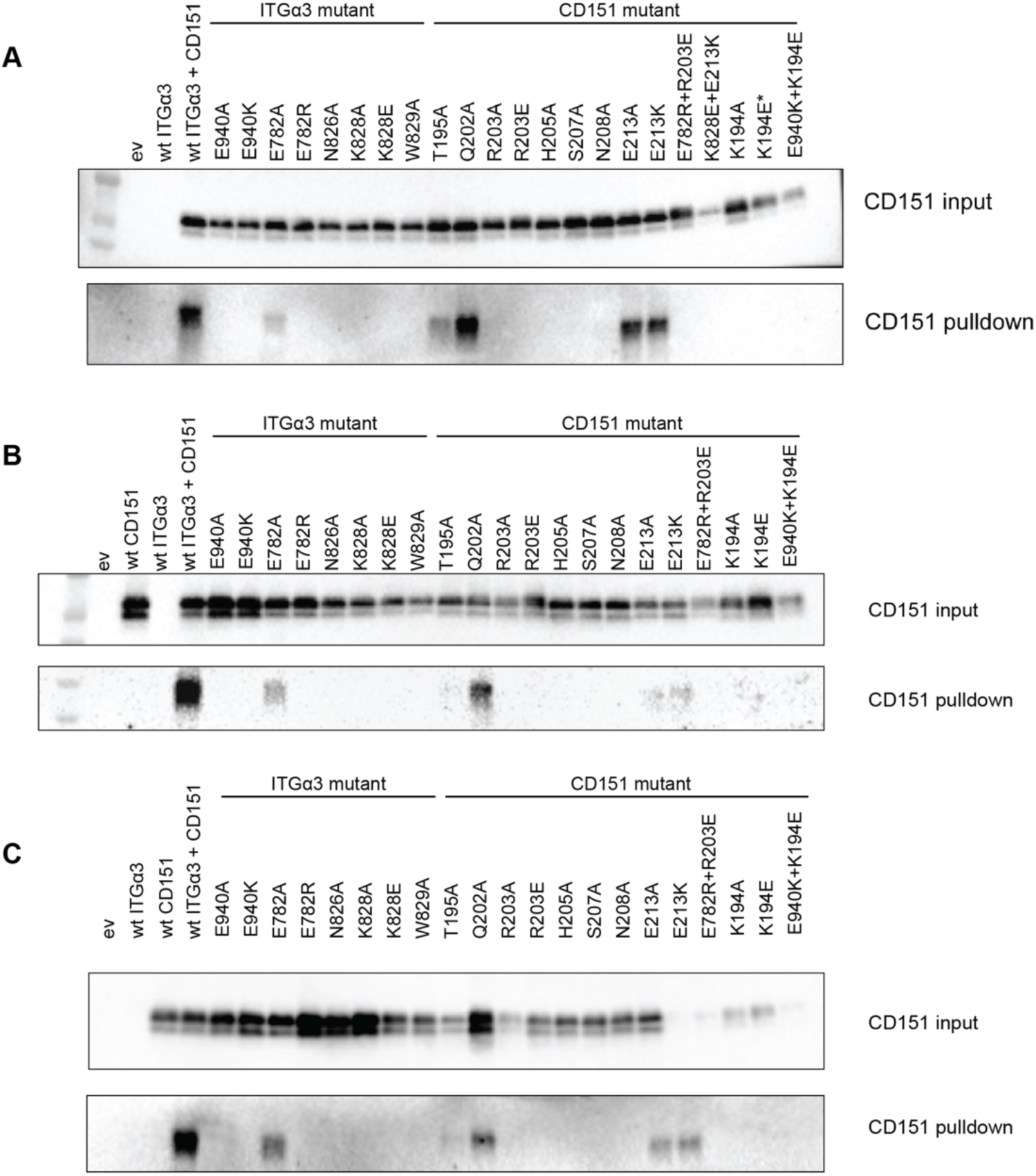
Effect of mutations at the ITGα3-CD151 interface on co-immunoprecipitation of CD151 and ITGα3. (A-C) SVG-A TKO cells were transfected with plasmids expressing the indicated CD151 and ITGα3 proteins, and complexation of CD151 with protein-C tagged ITGα3 was analyzed by co-immunoprecipitation using an anti-protein C antibody. The TS151 α-CD151 antibody was used for immunoblotting. Input represents 5% of the solubilized membrane fraction. Immunoprecipitations in cells transfected with only ITGα3 or CD151 served as negative controls. n = 3 independent experiments are shown.

**Figure S7.**
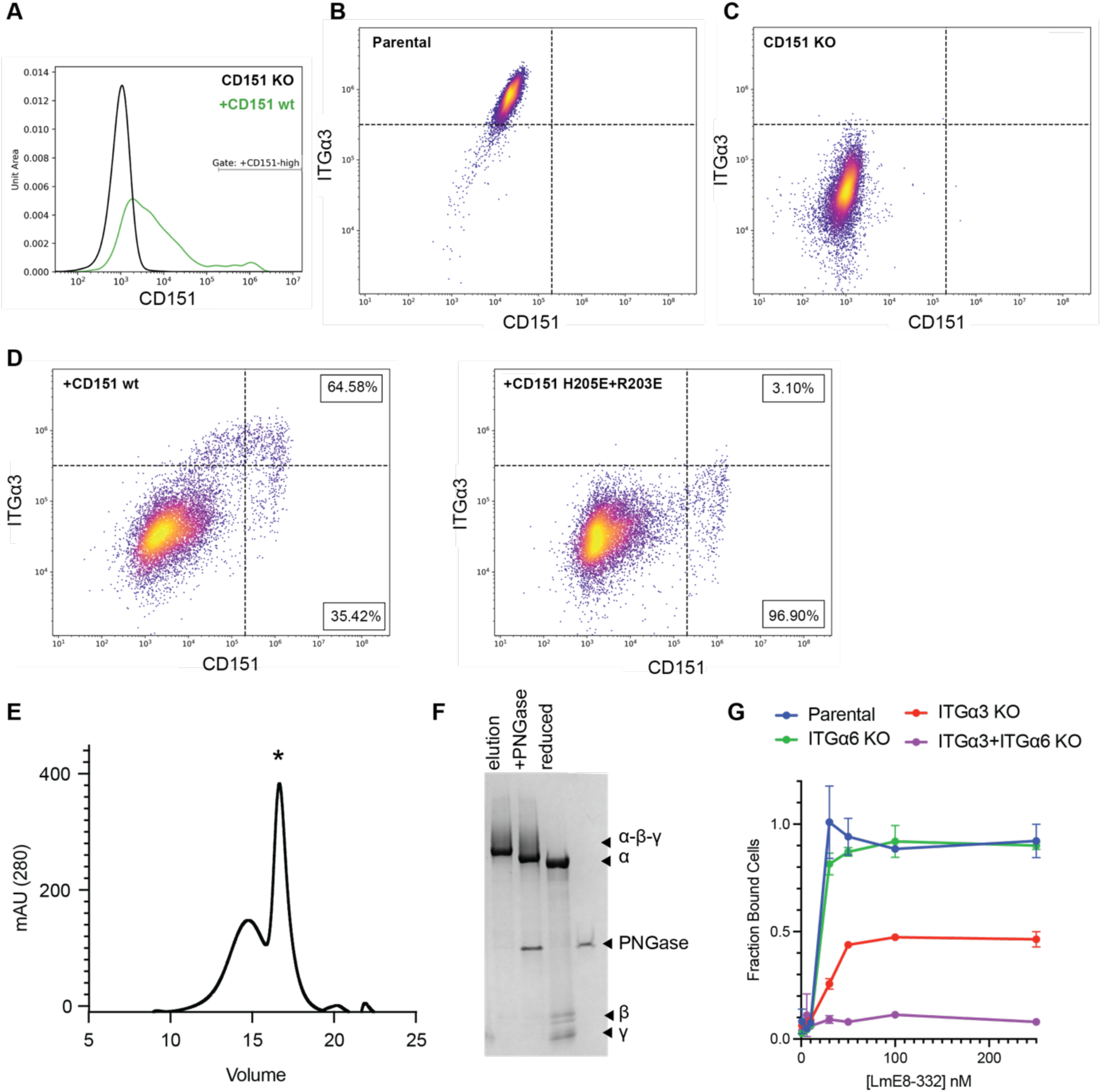
Flow cytometry gating strategy for analysis of CD151 rescue of ITGα3 surface export and purification of LmE8-332. (A) CD151 KO cells were transiently transfected with a plasmid expressing wild-type CD151, and cells with high CD151 expression were gated for analysis. This gating strategy was also used for surface export analysis after transfection with plasmids expressing CD151 mutant and tetraspanin-mNeonGreen fusion proteins. (B,C) Flow cytometry plot showing surface staining of ITGα3 as a function of CD151 expression in parental (B) and CD151 KO cells (C). Upper and lower quadrants distinguish between parental and CD151 KO surface staining for ITGα3. The right quadrant corresponds to the CD151 positive cells used to analyze rescue of ITGα3 surface staining. (D) Flow cytometry plot showing surface staining of ITGα3 as a function of rescue with either wild-type (left) or R203E/H205E mutant CD151 (right). (E-G) Purification and functional analysis of of LmE8-332. (E) Size exclusion chromatogram of LmE8-332 after affinity chromatography. (F) Coomassie stained gel showing LmE8-332 purity in the size exclusion peak indicated with an asterisk. Lanes showing deglycosylation by PNGase, and migration positions of the individual subunits (α, β, and γ) under reducing conditions are also shown. (G) Plot showing the effect of knocking out ITGα3 and ITGα6 on cellular adhesion as a function of LmE8-332 concentration coating the plates.

**Figure S8.**
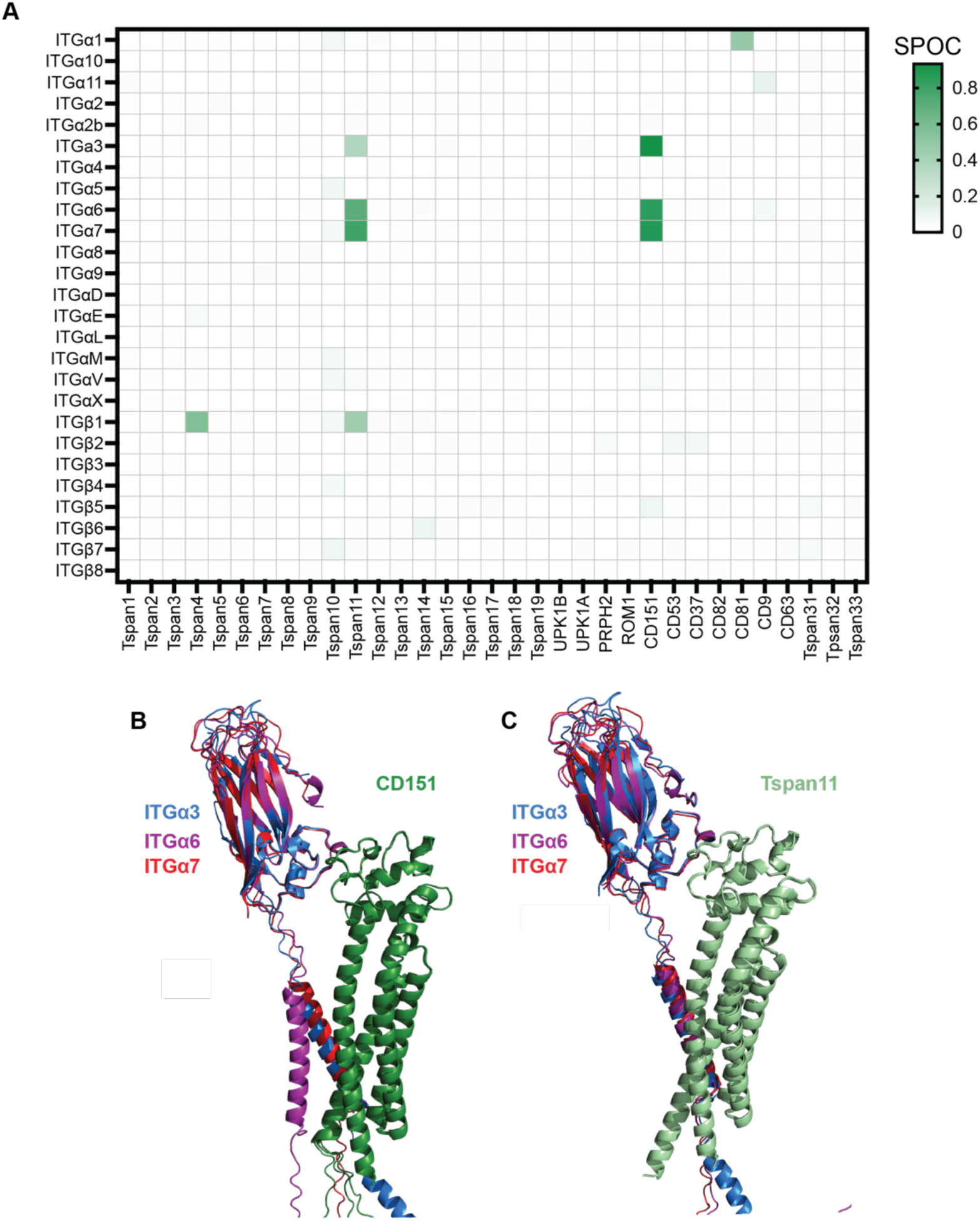
AlphaFold Multimer predictions of integrin-tetraspanin interactions. (A) Alpha Multimer predictions were evaluated using the Structure Prediction and Omics-Classifier (SPOC), which ranks the likelihood of a true interaction on a scale from 0 (low probability) to 1 (high probability, green). Only interactions of CD151 and Tspan11 with laminin binding integrins were confidently predicted. (B,C) Cartoon representation of Alphafold2 predictions of CD151 (B) and Tspan11 (C) complexes with the calf-2 and transmembrane regions of the ITGα3, ITGα6, and ITGα7. Note that the predicted interface is highly overlapping for all complexes of CD151 (B) and Tspan11 (C) with laminin binding integrins.

